# Engineering a performance-improved, axon-targeted kalium channelrhodopsin for optogenetic neuropathway inhibition

**DOI:** 10.64898/2026.01.17.700073

**Authors:** Simon Miguel M. Lopez, Han-Ying Wang, I-Chen Lee, Wei-Hsin Chen, Yong-Cyuan Chen, Yu-Jie Lin, Chien-Chang Chen, Ming-Kai Pan, Ching-Lung Hsu, Wan-Chen Lin

**Affiliations:** Taiwan International Graduate Program in Molecular Medicine, National Yang Ming Chiao Tung University and Academia Sinica, Taipei, Taiwan; Institute of Biomedical Sciences, Academia Sinica, Taipei, Taiwan; Department and Graduate Institute of Pharmacology, National Taiwan University College of Medicine, Taipei, Taiwan; Molecular Imaging Center, National Taiwan University, Taipei, Taiwan; Neuroscience Program of Academia Sinica, Academia Sinica, Taipei, Taiwan; Biomedical Translation Research Center, Academia Sinica, Taipei, Taiwan; Department of Life Science, National Taiwan University, Taipei, Taiwan

## Abstract

Optogenetic neuropathway inhibition is a powerful approach for dissecting circuit functions. This strategy, however, frequently encounters practical challenges due to insufficient expression or performance of the optogenetic silencer on axonal projections/terminals. HcKCR1, a light-gated potassium-selective channel from *Hyphochytrium catenoides*, has shown great promise for optogenetic inhibition. Unfortunately, the application of HcKCR1 in neuropathway manipulations is hindered by its unsatisfactory gating properties and poor axonal trafficking. To overcome these hurdles, we first engineered a performance-improved HcKCR1 (piKCR) that allowed more reliable neuronal inhibition at low intensities of green or red light. We next engineered an axon-targeted piKCR (piKCR.AT) that demonstrated long-range axonal trafficking and optical presynaptic inhibition in the mouse hippocampus. When piKCR.AT was expressed in the cerebellar Purkinje Cells (PCs), optical manipulation of PC outputs to the deep cerebellar nuclei robustly disrupted mouse movement on the balance beam. With enhanced performance and axonal distribution, piKCR.AT may provide new opportunities for elucidating neuropathway functions in health and diseases.

## Introduction

Optogenetics is a revolutionary technique that integrates optical stimulation and genetic engineering to precisely control biological activities in defined space, time, and cells (1–3). In neuroscience, opsins such as light-gated ion channels/pumps are frequently employed to optogenetically control neuronal activities. Light can be delivered to the somata of opsin-expressing cells to stimulate or suppress their firing, or to one of their projection pathways to manipulate signal delivery to a specific downstream target. To functionally dissect neurocircuits underlying a specific behavior or a physiological process, reliable silencing of a defined neuropathway is highly desirable. Such manipulation requires sufficient expression of an inhibitory actuator at the projection terminals, as well as powerful performance of the inhibitor to overcome the incoming action potentials (4).

Among the large variety of optogenetic silencers, prevailing options include light-driven ion pumps (e.g., NpHR (5) and ArchT (6)), anion-conducting channelrhodopsins (ACRs; e.g., GtACR1/2 (7)), and inhibitory optogenetic G-protein-coupled receptors (opto-GPCRs; e.g., eOPN3 (8) and PdCO (9)). Activation of light-driven ion pumps directly hyperpolarizes the membrane potential, offsetting the baseline for neuronal excitation. Due to one-to-one stoichiometry between photon absorption and ion pumping, this mechanism requires high opsin-expression level and bright illumination to achieve robust silencing. These technical requirements, together with the active transport nature of ion pumps, could lead to adverse effects such as tissue heating (10), pH change (11), or intracellular chloride accumulation (which may alter the reversal potential of the GABA_A_ receptor (12)). Chloride-conducting ACRs functionally resemble GABA_A_ receptors, inhibiting neuronal firing through not only hyperpolarization but also shunting (2, 4). However, activation of ACRs in axons/presynaptic terminals may result in membrane depolarization and even neuronal excitation due to reduced transmembrane chloride gradients in these compartments (2, 4, 11, 13). Opto-GPCRs have demonstrated powerful presynaptic inhibition in both brain slice electrophysiology and *in vivo* experiments (8, 9). However, activation of opto-GPCRs may stimulate multiple G-protein coupled signaling pathways that affect not only neuronal excitability but also other cellular/subcellular events (2). The engagement of multiplexed effector responses might compromise the temporal precision and/or complicate the causal relationship of the optical manipulation through opto-GPCRs.

In the nervous system, potassium channels play critical roles in terminating action potentials or setting neuronal excitability. Light-driven actuators that functionally mimic endogenous potassium channels are thus long-sought optogenetic inhibitors. Natural potassium-selective channelrhodopsins (i.e., kalium channelrhodopsins; KCRs) from *Hyphochytrium catenoides* (HcKCR1/2) (14) and *Wobblia lunata* (WiChR) (15) were recently discovered as promising silencers that exhibit exceptionally high sensitivity to light. Several variants of KCR have been applied in small model organisms (16, 17) and mice (15, 18, 19) to demonstrate potent neuronal inhibition *in vivo*. Notably, a new variant of HcKCR1 (HcKCR1-hs), was employed in rodent epilepsy models to achieve transcranial optogenetic seizure suppression (19). Despite these promising observations, targeted neuropathway inhibition by KCRs has not been reported yet. The paucity may be largely attributed to the inefficient axonal trafficking of KCRs. As reported for HcKCR1-hs (whose cell-surface expression was maximized by its N-terminal signal peptide, plasma membrane trafficking signal, and ER export motif), the opsin was not noticeable in axons when expressed in the hippocampal pyramidal neurons (19). Additionally, unsatisfactory functional properties such as desensitization (14) or ion selectivity shift under prolonged illumination (14, 17, 20) might further impose constraints to KCRs for mediating neuropathway inhibition *in vivo*.

Here, we report a systematic engineering of HcKCR1 for optogenetic neuropathway manipulation. We found that substitution of T109, a residue near the retinal binding site, with bulkier hydrophobic amino acids can alleviate desensitization and enhance stationary current under continuous illumination. By mutating T109 and H225 (a residue that influences the channel’s potassium selectivity; Ref. 18) together, we engineered a **<u>p</u>**erformance-**i**mproved Hc**<u>KCR</u>**1 (piKCR) that enabled efficient action potential suppression under low levels of green or red light. We further developed an axon-targeted piKCR (piKCR.AT) which distributed efficiently in long-range axonal projections and enabled optogenetic presynaptic inhibition. When expressing piKCR.AT in the cerebellar Purkinje cells (PCs) of mice, optically manipulating PC outputs to the deep cerebellar nuclei (DCN) caused significant movement disruption. With its robust inhibitory effect and successful axonal expression, piKCR.AT serves as a potent optogenetic silencer for probing neuropathway functions in health and diseases.

## Results

### Engineering a performance-improved kalium channelrhodopsin (piKCR) through structure-guided mutagenesis

Reliable optogenetic inhibition requires robust and sustained opsin activity over the course of manipulation. However, several studies have reported that HcKCR1 desensitizes rapidly (14, 15) and exhibits a depolarizing shift of reversal potential (which reflects the decrease of the channel’s potassium selectivity; Refs. 14, 15, 17, 21) under prolonged illumination. To achieve reliable neuropathway inhibition, we first explored mutations in HcKCR1 that allow systematic improvement of its gating properties. In HcKCR1, retinal forms a Schiff base with K233 (Fig. 1A), and its photoisomerization triggers protein conformational changes that lead to the formation of the potassium-conducting pathway (18, 22). T109, a residue that is close to the retinylidene Schiff base and involved in the light-induced conformational rearrangements (Fig. 1A) (18, 22), appears to be a candidate site for modifying HcKCR1 gating. Two previously reported mutants, T109A (18) and T109V (24), exhibited altered rates in channel deactivation or desensitization. We therefore prepared a series of T109 mutants and compared their photocurrent properties with those of the wild-type HcKCR1 measured from human embryonic kidney (HEK) cells under a bi-ionic recording condition (Fig. 1B–H). Each tested opsin was fused with a C-terminal enhanced yellow fluorescent protein (EYFP) for visualization in HEK cells.

**Fig. 1.**
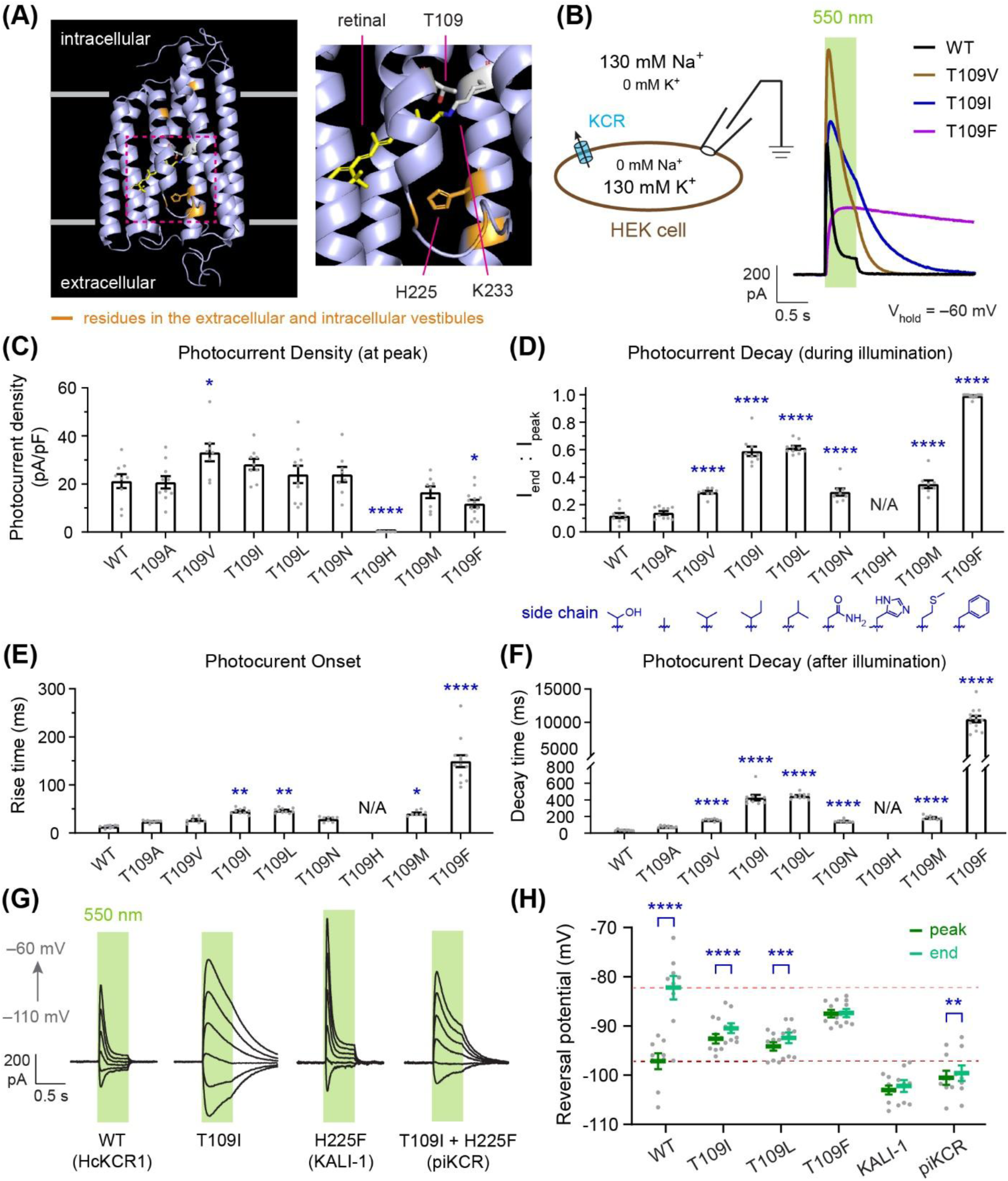
Engineering a performance-improved kalium channelrhodopsin (piKCR). (**A**) The location of amino acid residue T109 in the dark state of HcKCR1 (PDB code: 8H86; Ref. 18) showing its close proximity to the retinylidene Schiff base (formed between the ε-amino group of K233 and the aldehyde group of retinal. Some key residues, including H225 (whose mutations may enhance the channel’s potassium selectivity; Ref. 18), in the extracellular and intracellular vestibules are colored in orange. (**B**) Representative voltage-clamp recording traces from HEK cells showing light-elicited currents from the wild-type HcKCR1 and T109 mutants under a bi-ionic condition (130 mM extracellular Na^+^ and 130 mM intracellular K^+^). (**C**–**F**) Comparison of the wild-type HcKCR1 with T109 mutants in their photocurrent density (**C**), end-to-peak current ratio (**D**, to assess the degree of desensitization), current onset time (**E**, measured as 10–90% rise time), and current decay time (**F**). N/A, not available due to the lack of measurable photocurrents. I_end_: photocurrent amplitude measured at the end of the illumination. I_peak_: amplitude of the photocurrent peak. The chemical structure of the amino acid side chain at position 109 is plotted for each tested construct in panel **D**. Cells were held at –60 mV. n = 8–13 cells. \**P* <0.05, \*\**P* <0.01, \*\*\*\**P* <0.0001 (one-way ANOVA with Dunnett’s multiple comparison’s test except for T109F in panel **F**, which was compared with the wild-type using unpaired t test). (**G**, **H**) Reversal potentials of the wild-type HcKCR1, the slow-desensitizing T109 mutants, KALI-1 (i.e., H225F mutant which exhibits higher potassium selectivity; Ref. 18), and piKCR. (**G**) Representative voltage-clamp recording traces showing photocurrents elicited at different holding potentials. (**H**) Reversal potential group data (n = 8–9 cells). Dark- and light-red dashed lines indicate the average reversal potentials at the current peak and the end of illumination, respectively, of the wild-type HcKCR1. \*\**P* <0.01, \*\*\**P* <0.001, \*\*\*\**P* <0.0001 (paired t test). The reversal potentials were measured from the current-voltage relationships of the recorded cells (Fig. S1). Additional statistics are listed in Table S1. For panels **C**–**H**, cells were illuminated with 550 nm (0.084 mW/mm^2^, 500 ms) to evoke currents. Recording were carried out using a bi-ionic condition shown in panel **B**. Data are presented as mean ± SEM, with individual data points shown as grey symbols.

All mutants except T109H produced appreciable outward currents in response to a 550-nm light flash (peak photocurrent density >10 pA/pF at –60 mV; Fig. 1, B and C). As reported previously (14, 15), the wild-type HcKCR1 desensitized quickly, resulting in only 10% current (compared to the peak amplitude) remained at the end of a 500-ms continuous illumination (Fig. 1B). To evaluate the effect of T109 mutation on desensitization, we measured each variant’s current amplitude at the peak (I_peak_) versus at the end of the illumination (I_end_). By comparing the end-to-peak ratio (I_end_ : I_peak_), we found that replacing T109 of the wild-type with a hydrophobic residue (e.g., V, I, L, and F) significantly alleviated desensitization and increased the sustained current (i.e., I_end_; Fig. 1, B and D). Intriguingly, the end-to-peak ratio appeared to elevate with the increasing size of the hydrophobic side chain: T109A (0.14 ± 0.01) < T109V (0.29 ± 0.01) < T109I (0.59 ± 0.03) ≈ T109L (0.61 ± 0.01) < T109F (0.99 ± 0.00) (Fig. 1D). Moreover, mutants carrying a polar residue (e.g., T or N) at position 109 desensitized more rapidly than mutants with a similarly sized hydrophobic residue (e.g., T→V or N→L, respectively; *P* < 0.0001 by unpaired t test; Fig. 1D).

Increasing the size of hydrophobic residue at position 109 decelerated not only desensitization (Fig. 1D) but also activation (Fig. 1E) and deactivation (Fig. 1F). The effect of mutation was more profound on deactivation than activation. For example, substituting T109 with isoleucine caused a 3.4-fold increase in the onset time (measured as 10-90% rise time) but a 12.5-fold increase in the post-illumination decay time of the photocurrent (wild-type: 13.4 ± 1.0 ms onset time, 34.3 ± 2.5 ms decay time; T109I: 45.4 ± 2.0 ms onset time, 427 ± 34 ms decay time; Fig. 1, E and F). Notably, the T109F mutant exhibited much slower desensitization and deactivation (0.99 ± 0.00 end-to-peak ratio and 10481 ± 477 ms decay time; Fig. 1, D and F), making it potentially suitable for long-timescale applications. Together, these results suggest that systematic tuning of HcKCR1 gating kinetics can be achieved by adjusting the size of hydrophobic residue at position 109.

We next addressed another potential issue of HcKCR1, that is, the depolarizing shift of reversal potential (E_rev_) during continuous illumination. Consistent with previous findings (14, 15, 21), we observed that E_rev_ of the wild-type HcKCR1 shifted from –97 ± 2 mV (at peak) to –82 ± 2 mV (at the end of the light flash) over the course of a 500-ms photostimulation (Fig. 1H and Fig. S1). This 15-mV E_rev_ shift (ΔE_rev_) reflects a decrease of the channel’s K^+^/Na^+^ permeability ratio (P_K_/P_Na_; calculated using a modified Goldman-Hodgkin-Katz voltage equation; Ref.14) from 38 ± 3 to 22 ± 2. Interestingly, the top three slow-desensitizing mutants (T109I, T109L, and T109F) did not exhibit such a rapid E_rev_ shift (Fig. 1H, Fig. S1, and Table S1). However, the reversal potentials of these mutants were somewhat less negative than the peak E_rev_ of the wild-type (especially T109F: peak E_rev_ = –88 ± 1 mV, *P* < 0.0001, one-way ANOVA; Fig. 1H and Table S1). Considering the channel’s potassium selectivity and temporal precision for optogenetic control, we chose T109I for further optimization. We employed a second mutation, H225F, which was previously reported to increase the potassium selectivity of HcKCR1 (18) (Fig. 1H and Fig. S1). Compared to the wild-type, the resulting double mutant (T109I + H225F) showed comparable E_rev_ at the photocurrent peak and very small ΔE_rev_ at the end of the continuous illumination (Fig. 1H, Fig. S1, and Table S1). Moreover, the double mutant elicited comparable photocurrent magnitude with the wild-type (Fig. S2A) and exhibited similarly slow desensitization as the T109I mutant (Fig. S2B). This new variant, which is named piKCR (**<u>p</u>**erformance-**i**mproved Hc**<u>KCR</u>**1), will serve as the template for subsequent functionalization.

### Characterizing a membrane-targeted piKCR (piKCR.TE) in HEK cells

EYFP-tagged HcKCR1 is known to suffer from incomplete membrane trafficking and expresses largely as intracellular aggregates (14), limiting its *in vivo* applications. Likewise, EYFP-tagged piKCR aggregated severely inside cells (Fig. 2A). This problem was significantly improved by incorporating a membrane-targeting sequence (TS) and an ER-export motif (EE) into piKCR to promote its trafficking (5) (Fig. 2A). The resulting membrane-targeted piKCR (piKCR.TE) was further characterized by electrophysiology using more physiologically relevant extracellular and intracellular solutions (Fig. 2B). We first confirmed that piKCR remained slower in desensitization than HcKCR1 after adding TS and EE motifs. As shown in Fig. 2B, the end-to-peak current ratio of the membrane-targeted HcKCR1 (HcKCR1.TE) was only 0.08 ± 0.06 under 525 nm illumination, whereas the end-to-peak current ratio of piKCR.TE was 5.5-fold higher (0.41 ± 0.02). We next measured the reversal potential of piKCR.TE at the photocurrent peak and at the end of the 525 nm illumination (–68 ± 1 mV and –67 ± 1 mV, respectively; Fig. 2B), observing only ∼1 mV drift during photostimulation (one second at 0.18 mW/mm^2^). These results indicated that mutations (T109I and H225F) enhancing the stationary current and stabilizing E_rev_ are compatible with the intracellular motifs that promote the opsin’s cell-surface expression.

**Fig. 2.**
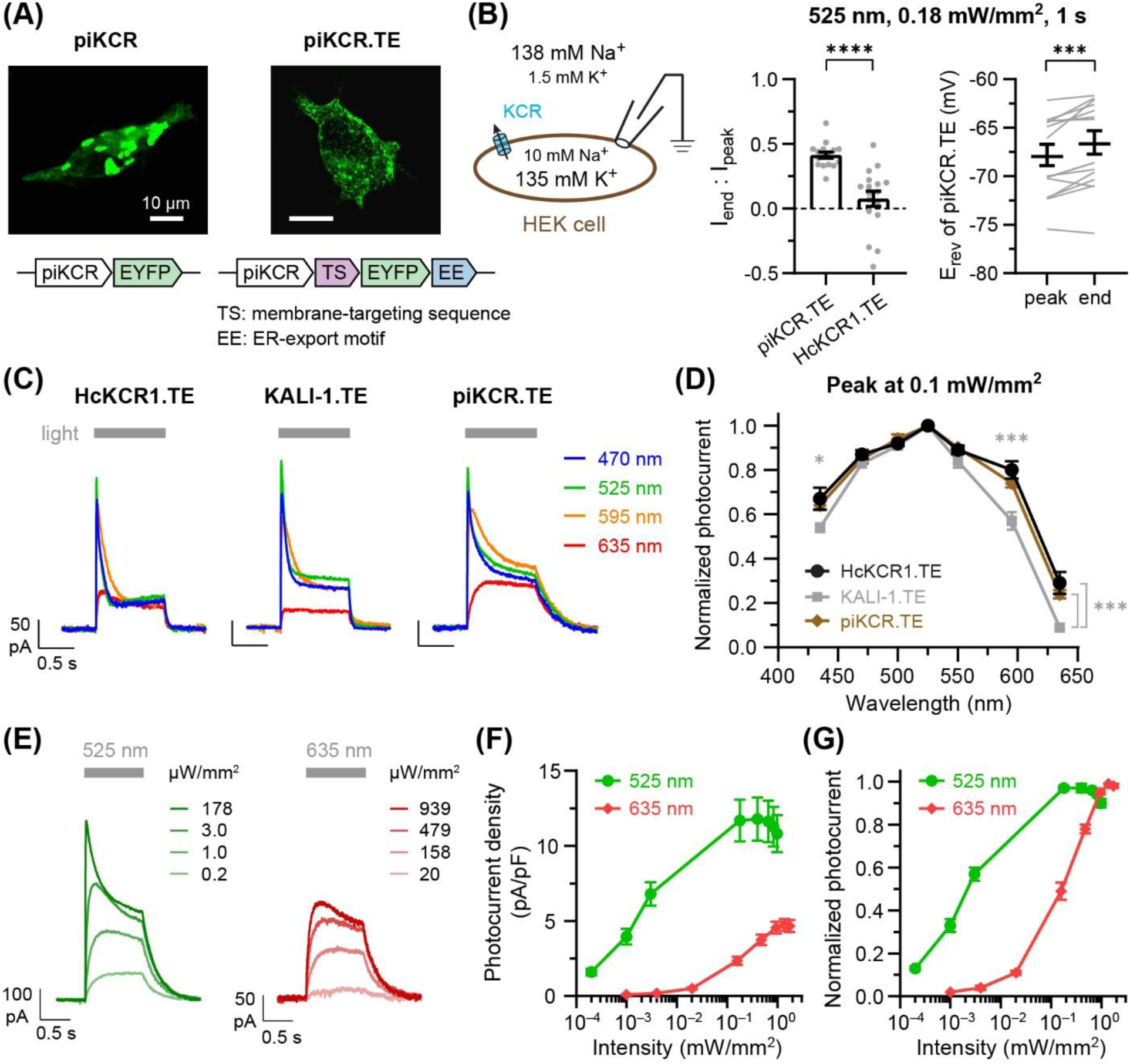
Photo-activation profiles of a membrane-targeted piKCR (piKCR.TE) in HEK cells. (**A**) Confocal images of HEK cells expressing piKCR or piKCR.TE demonstrating reduction of intracellular opsin aggregation by the incorporation of TS and EE motifs. Opsin localization is indicated by the green color (i.e., fluorescence from the fused EYFP). (**B**) Electrophysiological characterization of piKCR.TE using physiological extracellular and intracellular recording solutions. Cells were held at held at –60 mV for measuring the end-to-peak current ratio. I_end_: photocurrent amplitude measured at the end of one-second 525 nm illumination. I_peak_: amplitude of the photocurrent peak. (*Left*) piKCR (n = 16 cells) remained to desensitize more slowly than HcKCR1 (n = 17 cells) after incorporating TS and EE motifs. \*\*\*\**P* < 0.0001, Welch’s test. (*Right*) The reversal potential (E_rev_) of piKCR.TE only exhibited a small depolarizing drift over one second of illumination (1.3 ± 0.3 mV; n = 16 cells). \*\*\**P* < 0.001, paired t test. (**C**, **D**) Action spectra of the membrane-targeted HcKCR1, KALI-1, and piKCR. (**C**) Representative photocurrent traces elicited by 0.1 mW/mm^2^ of blue, green, orange, and red light. (**D**) Action spectra (n = 6, 9, and 7 cells for HcKCR1.TE, KALI-1.TE, and piKCR.TE, respectively). \**P* <0.05, \*\*\**P* <0.001 (for KALI-1.TE; one-way ANOVA with Dunnett’s multiple comparison’s test). (**E**–**G**) Light sensitivity profiles of piKCR.TE stimulated by green or red light (n = 13 and 10 cells, respectively). (**E**) Representative photocurrent traces. (**F**) Light sensitivity profiles plotted as peak photocurrent density vs. light intensity. (**G**) Light sensitivity profiles plotted as normalized photocurrent peak vs. light intensity. Action spectra and light sensitivity profiles were measured from HEK cells (held at –60 mV) using the physiological ionic conditions shown in panel **B**. Group data in panels **B**, **D**, **F**, **G** are plotted as mean ± SEM. Individual measurements in panel **B** are shown as grey symbols (I_end_ : I_peak_) or grey lines (E_rev_).

We further examined whether the mutations affect the optical properties of the opsin. The action spectrum of piKCR.TE (at 0.1 mW/mm^2^; Fig. 2, C and D) was compared with those of HcKCR1.TE and KALI-1.TE (i.e., membrane-targeted H225F mutant). As reported previously (18), H225F mutation significantly reduced the opsin’s responsiveness to long wavelengths of light (595 nm and 635 nm; KALI-1.TE in Fig. 2D). Although piKCR.TE also carried the H225F mutation, its action spectrum was nearly identical to that of HcKCR1.TE (Fig. 2D), eliciting maximal current (I_max_) at 525 nm, producing robust activities (>70% of I_max_) with 470–595 nm light, and responding to 635 nm moderately (achieving ∼25% of I_max_). The dynamic range of piKCR.TE activation was determined by its light sensitivity profile (Fig. 2, E–G). piKCR.TE responded to green light at very low intensities (achieving 13 ± 1% I_max_ at 0.2 μW/mm^2^) and reached maximal activation at ∼200 μW/mm^2^ (Fig. 2G). Red light was effective but less potent than green light: photostimulation of piKCR.TE required 20 μW/mm^2^ and 1.4 mW/mm^2^ of 635 nm light to reach ∼10% and maximal activation, respectively (Fig. 2G). Moreover, I_max_ evoked by 635 nm was ∼40% of I_max_ by 525 nm (Fig. 2F). Notably, however, red light produced stable currents at most tested intensities, while green light caused more profound desensitization at >1 μW/mm^2^ (Fig. 2E). The enhanced stationary photocurrent, stable E_rev_, and substantial responses to green and red light at the μW/mm^2^ level make piKCR.TE potentially suitable for *in vivo* applications.

### piKCR.TE efficiently suppresses neuronal firing through gain control

After characterizing basic properties of piKCR.TE in HEK cells, we evaluated its performance using whole-cell current clamp recordings in cultured cortical neurons (Fig. 3). The opsin distributed extensively in neurons (Fig. 3A) and suppressed action potentials rapidly and reversibly upon light switching (Fig. 3B). We first determined the minimal light intensity required to eliminate action potentials within a train of spikes elicited by continuous injection of a depolarizing current (100-150 pA; Fig. 3, B and C). Green light (525 nm) was extremely potent for piKCR.TE-expressing cells, hyperpolarizing membrane potential and eliminating spikes at the lowest intensity tested (0.2 μW/mm^2^; Fig. 3, B and C). Consistent with the action spectrum and photosensitivity profiles of piKCR.TE in HEK cells (Fig. 2, D, F, and G), red light (635 nm) was less potent than green light; however, it also efficiently suppressed spikes at an appreciably low intensity (20 μW/mm^2^; Fig. 3, B and C). As shown in HEK-cell recordings (Fig. 2E), low intensities of green (<1 μW/mm^2^) and red (≤0.2 mW/mm^2^) light allow piKCR.TE to elicit steady currents during continuous illumination. To test whether such low levels of light are sufficient for reducing excitability of quiescent neurons, we delivered green or red light prior to injecting depolarizing currents (Fig. 3, D and E). The resting potentials of the recorded neurons were –62 ± 1 mV in the dark. Because the reversal potential of piKCR.TE was –68 ± 1 mV under physiological ionic conditions (Fig. 2B), light-mediated hyperpolarization was small (7.0 ± 1.1 and 5.5 ± 1.0 mV by 0.2 μW/mm^2^ 525 nm and 20 μW /mm^2^ 635 nm, respectively; Fig. 3, D and E). Surprisingly, both green and red lights strongly prevented neuronal spiking upon current injections up to +300 pA (Fig. 3, D and E). The strong reduction of neuronal excitability by red light is a unique feature of piKCR.TE, as HcKCR1.TE (which expressed similarly as piKCR.TE; Fig. 3A) did not prevent action potentials as significantly as piKCR.TE using the same experimental settings (Fig. 3F). Moreover, piKCR.TE enabled spiking to be suppressed within milliseconds using <1 μW/mm^2^ green light (or <50 μW/mm^2^ orange light), allowing optical inhibition to be achieved with good temporal precision (Fig. S3).

**Fig. 3.**
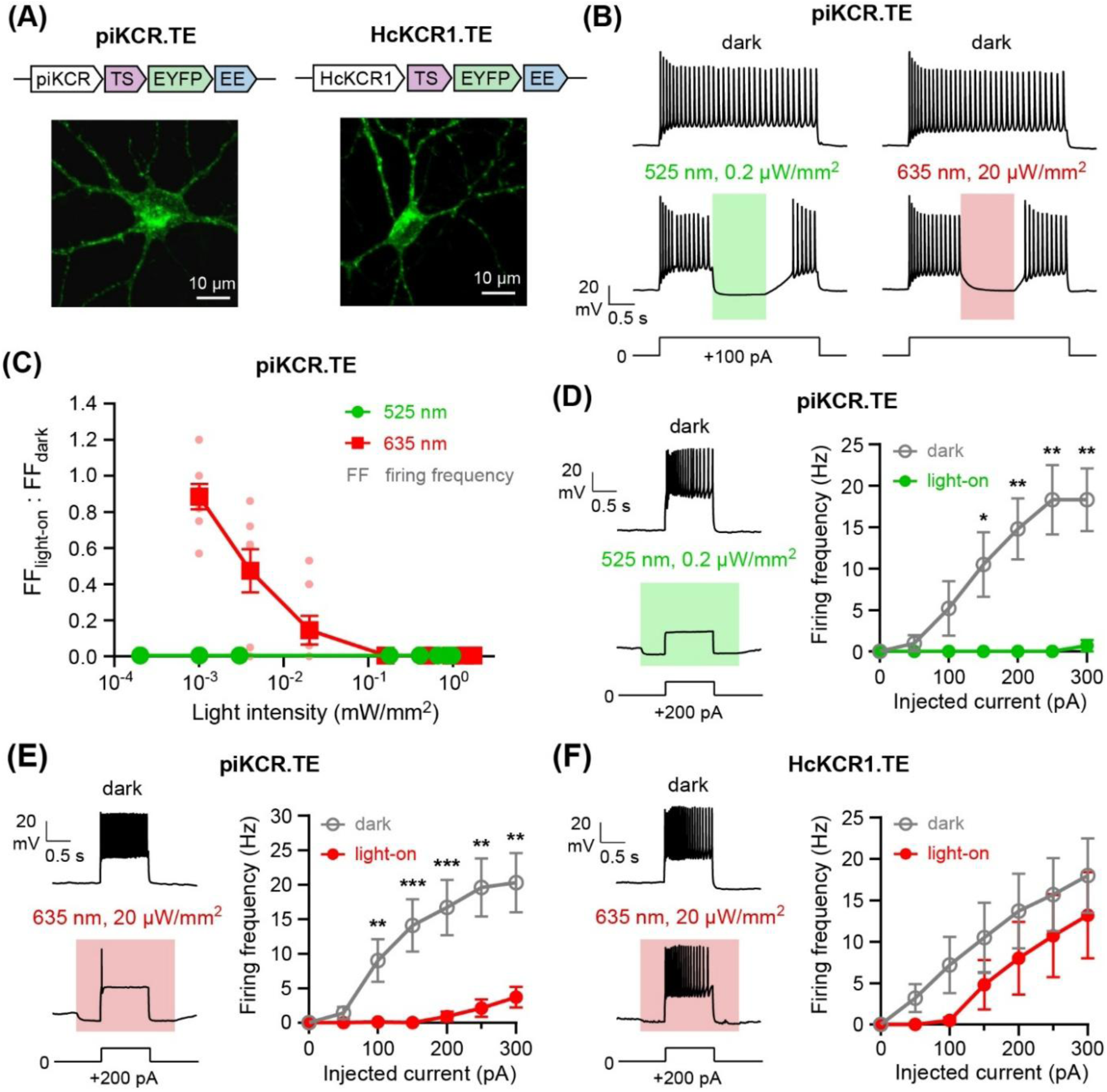
piKCR.TE efficiently suppresses neuronal spiking at low levels of light. (**A**) Confocal images of cultured cortical neurons expressing piKCR.TE or HcKCR1.TE. (**B**) Representative current-clamp recording traces from neurons expressing piKCR.TE, demonstrating reversible light-mediated spike suppression. In each trace, a train of action potentials were elicited by injecting 3 seconds of +100 pA current, with or without 1-second illumination in the middle of current injection. (**C**) The degree of action potential suppression (defined as firing frequency during illumination vs. in the dark) measured at different light intensities, using the experimental protocol illustrated in panel **B**. n = 5 and 7 cells for 525 nm and 635 nm, respectively. (**D**–**F**) Comparison of firing frequencies (elicited by injecting different amounts of depolarizing current) in KCR-expressing neurons with or without light conditioning. For recordings with light conditioning, illumination was started and terminated at 500 ms before and after current injection, respectively. (**D**) piKCR.TE: 525 nm, 0.2 μW/mm^2^; n = 6 cells. (**E**) piKCR.TE: 635 nm, 20 μW/mm^2^; n = 7 cells. (**F**) HcKCR1.TE: 635 nm, 20 μW/mm^2^; n = 6 cells. \**P* <0.05; \*\**P* <0.01; \*\*\**P* <0.001; Mann-Whitney test.

It is generally believed that light-activated ion pumps (e.g., eNpHR3.0 and eArch3.0) silence neurons by membrane hyperpolarization, while light-gated ion channels (e.g., GtACR1/2 or iC++) may “shunt” membrane depolarization if the channel’s reversal potential is close to the neuron’s resting potential (4, 25). Hyperpolarization and shunting are two distinct modes of neuronal inhibition: the former offsets/subtracts membrane depolarization but the latter reduces the neuron’s responsiveness (i.e., gain) to the excitatory stimuli. Small membrane hyperpolarization but strong spike prevention by piKCR.TE (Fig. 3, D and E) suggests that a “gain reduction” mechanism is involved in neuronal inhibition. This hypothesis was further supported by a light-mediated slope decrease in the subthreshold membrane potential versus current input relationship from piKCR.TE-expressing neurons (Fig. 4, A–C). The gain reduction mechanism may also allow piKCR.TE to silence neurons whose resting potentials are more negative than the opsin’s reversal potential. Indeed, when piKCR.TE was expressed in the anterior paraventricular thalamus (PVA) of mice, light weakly depolarized the neurons at rest (from –71 ± 1 mV before light to –65 ± 1 mV under illumination) but completely prevented action potential firing during current injection (Fig. 4, D and E). Together, our data demonstrate that piKCR.TE can robustly silence neurons without drastically altering the membrane potential. This feature could minimize the ionic flux and therefore prevent non-physiological changes in ionic concentrations during optical manipulations.

**Fig. 4.**
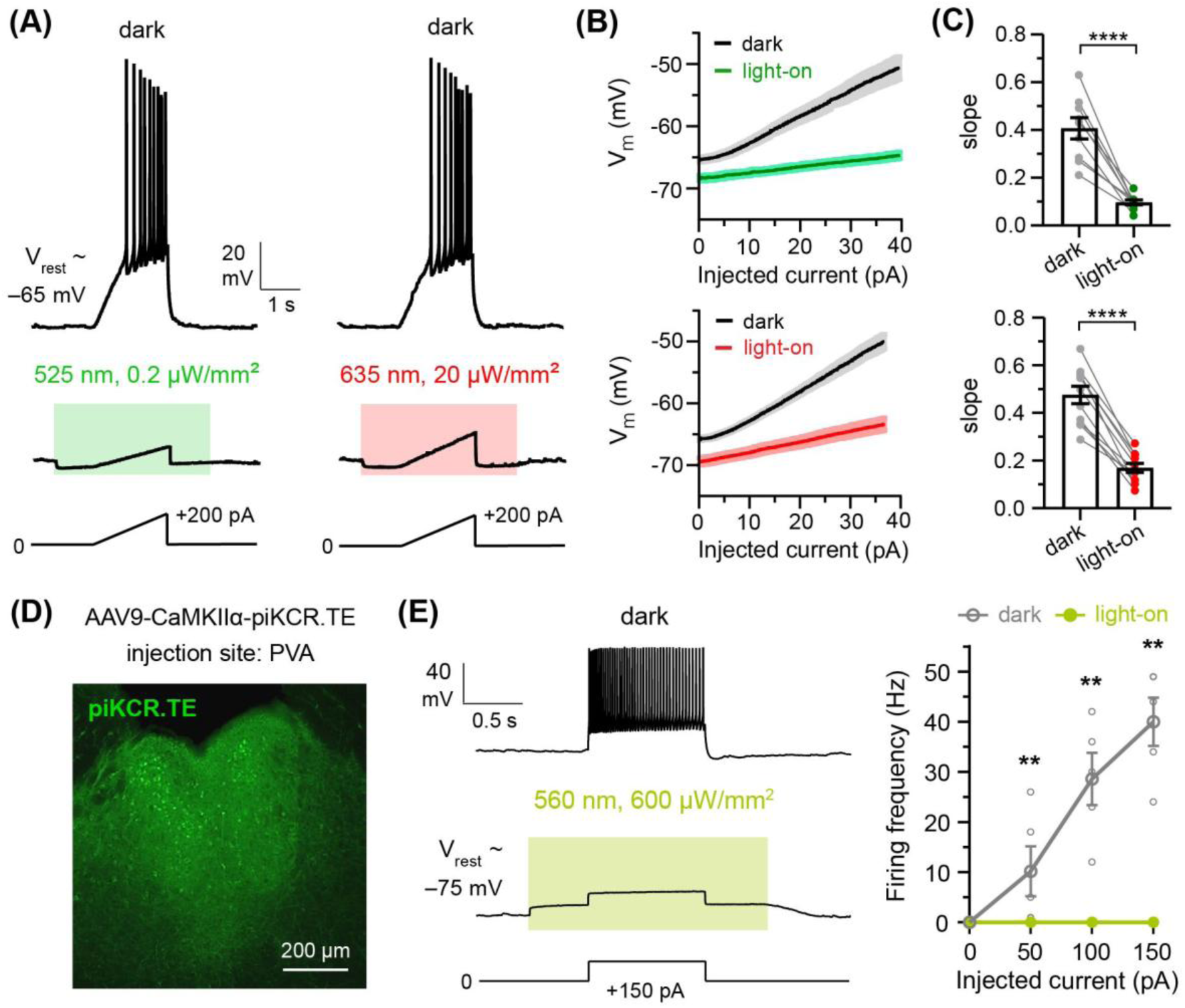
piKCR.TE can robustly prevent neuronal spiking without causing large membrane hyperpolarization. (**A**–**C**) piKCR.TE mediates optical reduction of neuronal gain. Current-clamp recordings of piKCR.TE-expressing cultured cortical neurons were carried out during 13–21 DIV (n = 9 and 11 cells for green and red light, respectively). Resting potentials of the recorded cells (initially ranged from –60 to –70 mV) were manually adjusted to –65 mV prior to the experiments. Depolarizing ramp current injections were applied with or without light conditioning. For recordings with light conditioning, illumination was started and terminated at 1 s before and after current injection, respectively. Green light (525 nm, 0.2 μW/mm^2^) and red light (635 nm, 20 μW/mm^2^) only caused a –3.8 ± 0.6 mV and –4.1 ± 0.6 mV change in membrane potential, respectively, prior to the ramp current injection. The slope of the subthreshold voltage-current relationship (i.e., the rate of membrane potential increase before the firing of the first spike) was significantly attenuated by both green and red light. (**A**) Representative recording traces. (**B**) The subthreshold voltage-current relationship plots (mean ± SEM) with or without light conditioning. (**C**) Group data (mean ± SEM) for subthreshold voltage-current slopes measured by linear fitting from individual neurons. \*\*\*\**P* <0.0001; Mann-Whitney test. (**D**, **E**) piKCR.TE enables optical neuronal silencing in brain slices. (**D**) A confocal image of mouse brain slice showing the expression of piKCR.TE in the anterior paraventricular thalamus (PVA). (**E**) Representative traces and group data (mean ± SEM, n = 5 cells) demonstrating piKCR.TE-mediated optical inhibition of spiking in PVA neurons. Note that action potentials were not elicited by current injections even though light induced a small baseline depolarization. \*\**P* <0.01; Mann-Whitney test.

### Engineering an axon-targeted piKCR (piKCR.AT) for long-range presynaptic inhibition

Robust neuropathway manipulation requires both high functional performance and successful axonal expression of the optogenetic tool. While HcKCR1 and its variants have exhibited powerful neuronal silencing *ex vivo* and *in vivo* (14, 16, 18, 19, 27), they have not been applied to neuropathway manipulation yet. This paucity is largely due to the poor trafficking and surface expression of HcKCR1 (14). Although surface expression of HcKCR1 can be strongly improved by incorporating an N-terminal signal peptide, a membrane trafficking signal (TS), and an ER export motif (EE) into the opsin, the distribution of this “enhanced” version is still restricted to soma and dendrites (19). These observations suggest that a specialized “axon-targeting” module is required for enabling HcKCR1 (and its mutants) to mediate neuropathway inhibition.

Several targeting strategies have been reported for delivering proteins to the axonal surface (28, 29). Considering HcKCR1’s inherited difficulty in cell-surface expression, we chose the intracellular domain (ICD) of neurexin (Nxrn) as its axon-targeting motif. The ICD of Nxrn (identical in both α and β isoforms) contains a PDZ-binding sequence that is responsible for the ER/Golgi exit, preferential expression on the axonal surface, and presynaptic targeting of Nxrn (30). It has been previously incorporated into a chemogenetic tool for selective presynaptic silencing (31). Likewise, we engineered an axon-targeted piKCR (piKCR.AT) by fusing the Nxrn ICD to the C-terminus of piKCR-EYFP (Fig. 5A).

**Fig. 5.**
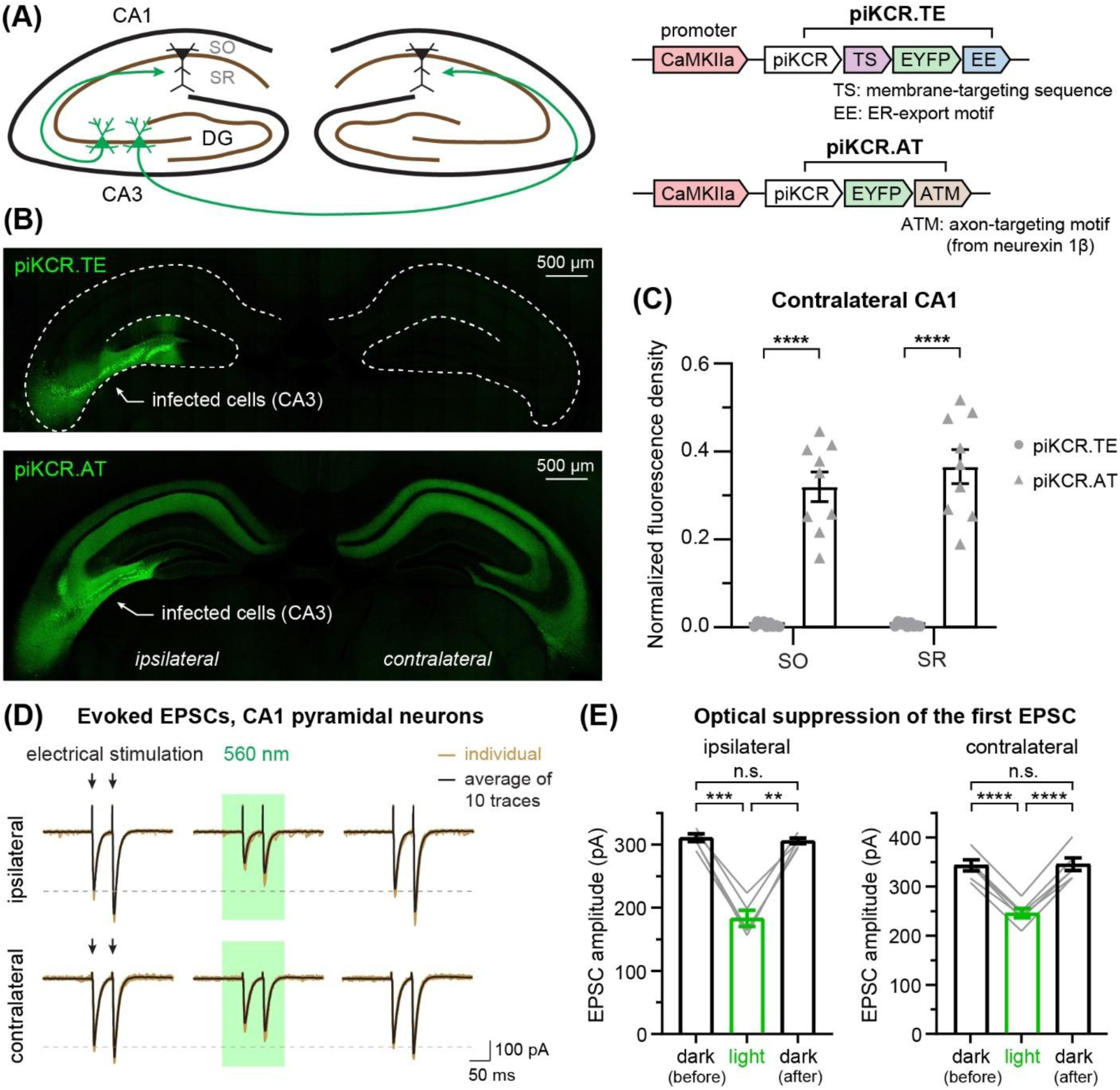
piKCR requires an axon-targeting motif to exert long-range presynaptic inhibition. (**A**) Schematic illustration showing unilateral expression of enhanced piKCR (piKCR.TE) or axon-targeted piKCR (piKCR.AT) in the mouse hippocampus. The infected neurons are shown in green, and the axonal projections of the CA3 pyramidal neurons (which may target CA1 neurons in either hemisphere) are shown as curved arrows. SO, *stratum oriens*; SR, *stratum radiatum*. (**B**) Confocal images of hippocampal slices (coronal sections) showing superior axonal expression of piKCR.AT (manifested as the fluorescence signals in CA1 SO or SR) to that of piKCR.TE. Slices were prepared from mice stereotaxically injected with AAV9 encoding either piKCR variant. (**C**) Comparing long-range axonal distribution of piKCR.TE vs. piKCR.AT using confocal images exemplified in panel **B**. The fluorescence densities of SO and SR layers in the contralateral CA1 subfield were normalized by the fluorescence density of the cell-body layer in ipsilateral CA3 (detailed in Methods). Group data are shown as mean ± SEM; n = 9 slices from 3 mice for both piKCR.TE and piKCR.AT. \*\*\*\**P* <0.0001 (Welch’s test). (**D, E**) piKCR.AT-mediated optical suppression of CA3→CA1 outputs. **D**, representative voltage-clamp recordings from near-horizontal hippocampal slices in which piKCR.AT was unilaterally expressed in CA3 pyramidal neurons (Fig. S4). CA1 pyramidal neurons were held at –70 mV, and excitatory postsynaptic currents (EPSCs) were electrically evoked by paired-pulse stimulation in the SR (with GABAergic activities blocked by gabazine). In both ipsilateral and contralateral CA1 neurons, the EPSC amplitudes were reduced by a 150-ms flash of green light (0.93 mW/mm^2^) and then fully recovered after light termination. **E**, group data from 5 (ipsilateral) and 6 (contralateral) cells shown as mean ± SEM. Measurements from individual cells are plotted as grey lines. \*\**P* <0.01; \*\*\**P* <0.001; \*\*\*\**P* <0.0001; n.s., not significant (repeated-measures one-way ANOVA with Geisser–Greenhouse correction followed by Tukey’s comparison).

To evaluate piKCR.AT in the mouse brain, we prepared an adeno-associated virus (AAV9) which encoded the piKCR.AT gene under a CaMKIIα promoter (Fig. 5A). Additionally, an AAV9 encoding piKCR.TE was prepared to test whether axonal delivery of piKCR is achievable by simply enhancing its surface expression. We examined axonal distribution of each opsin by injecting the respective virus to the hippocampal CA3 subfield unilaterally, followed by confocal imaging of brain sections 2–3 weeks after virus transduction. A CA3 pyramidal neuron may send axonal projections to the ipsilateral and/or contralateral CA1 regions (Schaffer-collateral and commissural pathways, respectively; Ref. 32). Accordingly, fluorescence signals (emitted from EYFP of the opsin) in CA1 would indicate axonal expression of the opsin in CA3 pyramidal neurons (Fig. 5A). Similar to the observation by Duan et al. (19), the distribution of piKCR.TE was restricted to ipsilateral CA3 but nearly undetectable in either CA1 region (Fig. 5B). In contrast, fluorescence from piKCR.AT was significant in not only ipsilateral CA3 but also CA1 regions of both hemispheres (Fig. 5B). The axonal levels of piKCR.TE and piKCR.AT were further compared using fluorescence density in the contralateral CA1 sublayers (normalized by the fluorescence density in the ipsilateral CA3 cell-body layer; Fig. 5C and Methods). In both *stratum oriens* (SO) and *stratum radiatum* (SR), the normalized fluorescence density of the piKCR.AT group is >50 times higher than that of the piKCR.TE group (Fig. 5C). These results suggest that the Nxrn ICD, when fused to the C-terminus of piKCR, can efficiently facilitate the trafficking of this opsin along both intra- and inter-hemispheric axonal projections.

We next electrophysiologically tested whether piKCR.AT enables optical inhibition of CA3→CA1 pathways in acute hippocampal slices (near-horizontal sections prepared from unilaterally injected mice described above; Fig. S4A). Excitatory postsynaptic currents (EPSCs) of ipsilateral or contralateral CA1 pyramidal neurons were evoked by paired-pulse stimulation delivered to the SR, with a stimulating electrode placed 200–300 μm away from the recorded cells. To test optical effects on presynaptic neurotransmitter release, a 150-ms flash of green light (0.93 mW/mm^2^ of 560 nm, started 50 ms prior to the first stimulation and ended after the second pulse) was delivered to the recorded region through the microscope objective (Fig. 5D). Light did not evoke currents prior to electrical stimulation (Fig. 5D) or alter the input resistance of the recorded cells (Fig. S4C), indicating no expression of postsynaptic piKCR. Compared to the baseline measurements in the dark, EPSC amplitudes were significantly reduced by green light (reduction in the first peak: ipsilateral, 41 ± 4%; contralateral, 29 ± 1%; Fig. 5, D and E). The reduced currents recovered back to the baseline levels after illumination, demonstrating the reversibility and speed of optical control (Fig. 5, D and E). It is important to note that the degree of EPSC reduction was constrained by the viral transduction efficiency in CA3 and the complexity of presynaptic inputs to the recorded CA1 neurons (see *Discussion*). Additionally, strong piKCR-mediated axonal inhibition could be implicated by the paired-pulse ratio (PPR): light did not cause a significant effect on PPR (Fig. S4B), suggesting that piKCR.AT likely eliminated action potentials before they could arrive at axonal terminals to trigger glutamate release (because light targeted both axonal projections and terminals). Light-mediated EPSC reduction was not observed in control experiments using mice injected with an EYFP-encoded AAV9 (Fig. S5), confirming that the inhibitory effect was contributed by piKCR.AT.

Expression of piKCR.AT (which carries a Nxrn ICD) in the presynaptic neurons did not significantly alter the baseline PPR: the ipsilateral PPR measured in the piKCR.AT group (1.32 ± 0.04, n = 5) was not different from that measured in the EYFP group (1.28 ± 0.02, n = 6; *P* = 0.4511, unpaired t test). Notably, the lack of light-evoked postsynaptic currents (Fig. 5D) suggested that activation of piKCR.AT in the CA3→CA1 axonal projections/terminals did not stimulate presynaptic neurotransmitter release. This observation highlights the advantage of piKCR.AT over ACRs, as axonal ACRs have been reported to stimulate rather than suppress neurotransmitter release (11, 13).

### Optical manipulation of the PC-to-DCN pathway in behaving mice

After validating axonal function of piKCR.AT in brain slices, we examined whether this tool allows optogenetic neuropathway manipulation *in vivo*. The cerebellum is well recognized for its role in enabling precise and coordinated movements (33). In the cerebellar cortex, Purkinje cells (PCs) serve as the principal neurons, sending inhibitory outputs to the deep cerebellar nuclei (DCN) to regulate signal integration and downstream outputs for learning, performing and fine-tuning motor actions (34–36). It is generally believed that PC dysfunction or degeneration, which weaken the inhibitory control over the DCN, may impair balance or motor coordination (36, 37). To directly assess functional impacts of the PC-to-DCN pathway, we expressed piKCR.AT in the PCs and tested whether illumination to the DCN could disrupt the motion of freely behaving mice (Fig. 6). An AAV9 encoding the gene of piKCR.AT was stereotaxically delivered to transduce neurons (including PCs) in the cerebellar cortex of the wild-type mice (Fig. 6, A and B). The presence of piKCR.AT (which carries an EYFP) in the PC projections/terminals was confirmed by observing fluorescent filaments/puncta in the DCN (Fig. 6B). To suppress the PC outputs, 589-nm light was illuminated onto DCN through an optical fiber. The effects of light on animal movement were measured using a balance beam walking assay (Fig. 6C). The mice were habituated and trained to conduct the walking task prior to testing light effects.

**Fig. 6.**
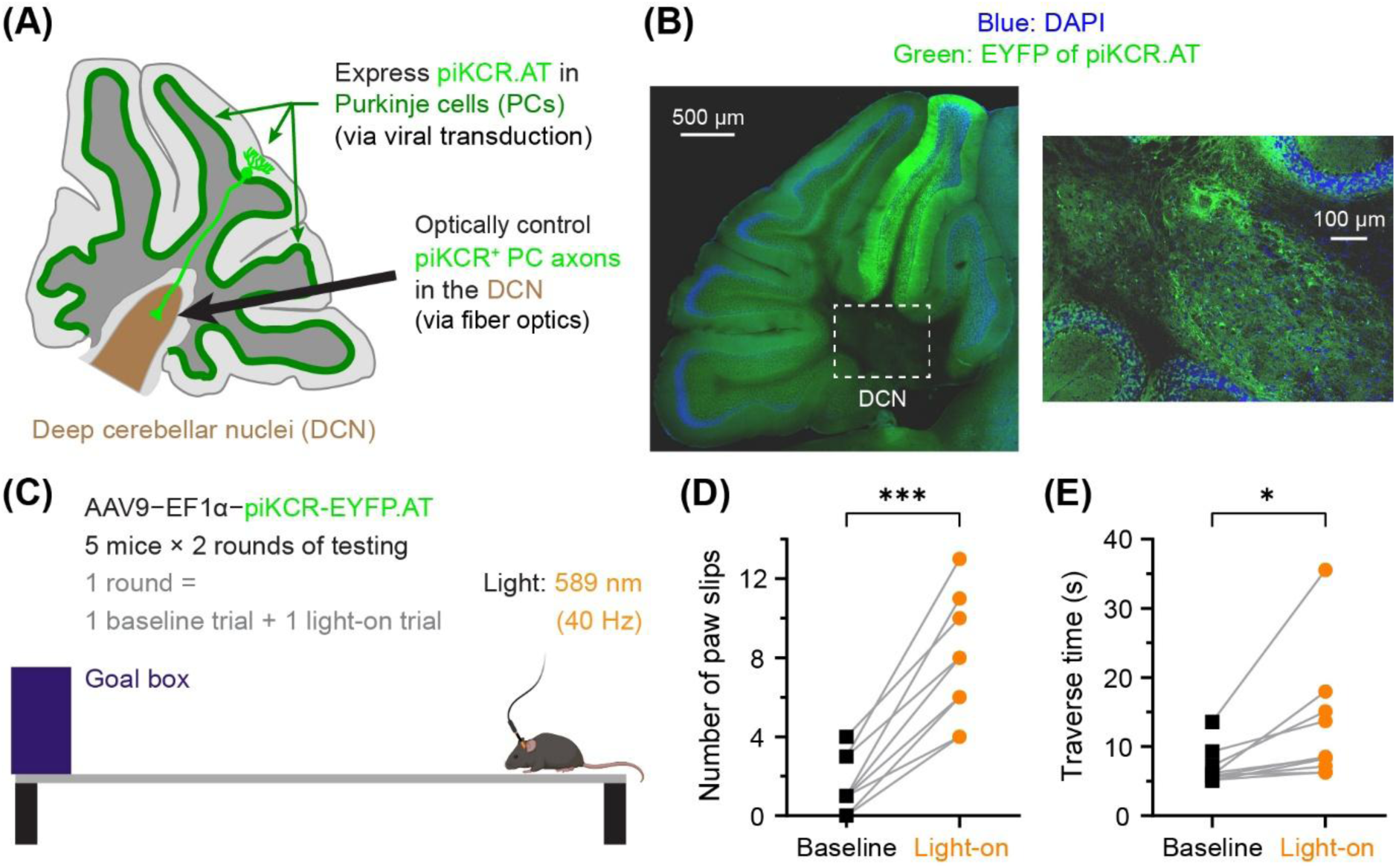
Optical manipulation of the PC-to-DCN pathway disrupted mouse movements on the balance beam. (**A**) A schematic illustration showing the location of a piKCR.AT-expressing Purkinje cell (PC; in green) sending its axon projection to the deep cerebellar nuclei (DCN; in brown). The PC layer is shown in dark green. In the behavioral experiment, the PC-to-DCN pathway was optically manipulated by delivering light to the DCN. (**B**) Confocal images of a sagittal cerebellar section from a mouse used in the behavioral assay. *Left*: a tiled image of the cerebellum cross-section. *Right*: an expanded image of the DCN (indicated by the dashed white box in the tiled image). (**C**) A schematic illustration for the balance beam walking assay. (**D**, **E**) Group comparison of walking behaviors showing disrupted mouse movements in the photocontrol (light-on) trials. (**D**) The number of foot slips during beam walking. (**E**) The amount of time used to complete beam walking. \**P* < 0.05, \*\*\**P* < 0.001; Mann-Whitney U test.

Under baseline (light-off) condition, the mice walked across the beam smoothly with only few occasions of foot slips (Fig. 6D and Movie S1). Illumination to DCN (which reduced the inhibitory inputs from PCs) disrupted the movements of the mice, significantly causing more foot slips and longer traverse time during beam walking (Fig. 6, D and E; Movie S2). This light-mediated movement disruption suggests that a sufficient population of piKCR.AT existed at the PC axonal terminals and robustly modulated PC outputs upon illumination. Moreover, these results demonstrate the significance of the PC-to-DCN projection in balance and motor coordination, providing a straightforward proof for this neuropathway’s physiological function.

## Discussion

Optogenetic neuropathway inhibition is a powerful approach for elucidating the operation mechanisms and/or physiological functions of a neural circuit. However, reliable tools for such manipulation are limited, and therefore the discovery of new silencers remains highly desirable. HcKCR1 has been considered as a promising candidate owing to its large conductance and high sensitivity to light. Although *in vivo* neuronal silencing by HcKCR1 and its variants have been achieved in several studies (16–20), targeted neuropathway inhibition by HcKCR1-based tools has been hindered by their poor axonal distribution. Moreover, the drifting of ion selectivity (14, 15, 21) and fast desensitization (14, 15) could make HcKCR1-mediated neuronal inhibition unreliable *in vivo* (17, 20). To overcome these limitations, we have strategically engineered an axon-targeted piKCR (piKCR.AT) that exhibited slow desensitization, stable ionic selectivity, and strong axonal distribution. With these improved properties, piKCR.AT may expand the repertoire of targeted neuropathway inhibition and open more opportunities for optogenetic applications.

For robust and reliable neuropathway inhibition in behaving animals, an opsin should provide substantial and stable inhibitory power over the course of optical manipulation. These criteria are manifested as slow desensitization and steady E_rev_ (i.e., stable potassium selectivity) for kalium channelrhodopsins. Although a few mutants have been found to decelerate channel closing and/or desensitization of HcKCR1 (18, 22–24), a systematic structure-function analysis of HcKCR1 gating kinetics remains pending. We here show that substituting T109, a residue that is near the retinylidene Schiff base and contributes to the central constriction site (18), with increasing sized hydrophobic amino acids (V, I/L, F) can progressively decelerate desensitization and channel closing of HcKCR1. Our observations indicate that hydrophobic interactions at/near position 109 appear to play a role in stabilizing the open state of HcKCR1, as mutants carrying a polar residue (e.g., T or N) desensitized/deactivated more rapidly than the similarly sized hydrophobic congener (V or L, respectively). Previous studies suggested that T109 may be involved in the light-induced conformational rearrangements that lead to channel formation of HcKCR1 (18, 22). Our findings support the unique role of T109 in HcKCR1 gating and provide practical information for tuning the kinetic properties of this opsin.

We engineered piKCR by combining a slow-desensitizing mutation (T109I) near the retinylidene Schiff base with a hyperpolarizing mutation (H225F) that stabilizes the architecture of the potassium-selectivity filter. Electrophysiological characterization of piKCR showed that the beneficial effects of individual mutations on channel gating/conduction were additive, and introducing both mutations to HcKCR1 did not significantly alter the opsin’s expression. Combination of mutations in distinct functional domains might thus be an effective approach for future piKCR design. For example, mutations at/near the retinal-binding site may combine with other hyperpolarizing mutations for the screening of new variants that exhibit enhanced sensitivity to red light, produce larger/more sustained stationary photocurrent, or open/close more rapidly.

Inhibition of neuronal spiking can be achieved by a subtractive mechanism (e.g., membrane hyperpolarization, which offsets the baseline neuronal excitation level) or a divisive mechanism (via gain control, i.e., reducing the neuron’s responsiveness to depolarizing inputs) (26). We found that piKCR.TE robustly prevents action potential firing from quiescent neurons in response to depolarizing input currents, even though light only causes a small hyperpolarizing shift of the membrane potential. The strong inhibitory effect arises from a significant reduction in the slope of the voltage output vs. current input relationship during illumination, suggesting that light exerts a divisive effect on the gain of piKCR-expressing neurons. Similarly, non-subtractive inhibitory effects were previously described for iC++, an engineered light-gated chloride channel (25). Berndt et al. compared neuronal inhibition efficiencies of eNpHR3.0 versus iC++ and found that eNpHR3.0 suppressed action potentials evoked by input currents whose magnitudes were equal to (or smaller than) that of the photocurrent. In contrast, iC++ could prevent spike generation in response to input currents that are much larger than the photocurrent it produced. Like piKCR.TE, iC++ did not cause strong hyperpolarization during illumination. Instead, the input resistance of iC++-expressing cells was strongly reduced by light. The authors suggested that efficient neuronal silencing by iC++ was likely caused by a mechanism similar to the shunting inhibition mediated by GABA_A_ receptors, the endogenous chloride-conducting channels responsible for neuronal inhibition.

Suppressing action potentials through light-gated ion pumps (mostly by light-mediated membrane hyperpolarization) *in vivo* may require high expression level of the opsins and/or strong illumination conditions to achieve robust effects. Such operations, however, might lead to unwanted ion accumulation or tissue heating. In this regard, piKCR.TE is advantageous owing to its high photosensitivity and strong reduction in neuronal responsiveness to depolarizing inputs. Because the reversal potential of piKCR.TE is close to the resting potential of most neurons, low intensity of light would cause only a small change to membrane potential but still strongly prevent action potential generation. The divisive nature of piKCR-mediated inhibition would therefore reduce the potential issues of ion accumulation and/or tissue heating over the course of *in vivo* manipulation. Moreover, piKCR.TE suppresses action potentials more robustly than HcKCR1.TE under red light (Fig. 3, E and F), making it an appealing tool for optogenetic neuronal inhibition in deeper tissues.

In this study, we demonstrated efficient axonal expression of piKCR.AT in two distinct types of neurons: the hippocampal CA3 pyramidal neurons and the cerebellar PCs. Appreciable axonal distribution of many opsins can be achieved by enhancing their expression on the neuronal surface (i.e., promoting their exit from ER and/or Golgi apparatus; Ref. 5). From our and others’ (19) observations, however, this approach is not effective for HcKCR1 variants. We found that piKCR requires a specialized motif to actively deliver itself to the axonal surface. The ICD of Nxrn turned out to be a successful axon-targeting module for piKCR, likely due to its multi-functional PDZ-binding motif that facilitates the ER/Golgi exit and polarized axonal transport of the opsin (30, 38). Moreover, it may stabilize the protein’s anchoring at the presynaptic terminal (39–41). It is worth noting that axonal transport of neurexin starts from trafficking of the newly synthesized protein to the dendritic surface, where it undergoes endocytosis, before transcytosis to the axon (38). Therefore, the distribution of piKCR.AT may be preferentially, but not exclusively, on the axon projections/terminals. In this regard, piKCR.AT would be more suitable for controlling neuropathway terminals that are spatially well segregated from their somatodendritic origin.

Both imaging and electrophysiology data suggested that piKCR.AT traveled successfully along the axons of CA3 pyramidal neurons, reaching ipsilateral and contralateral CA1 neurons to enable light-mediated presynaptic inhibition. It is worth noting that the effects we quantified from the patch-clamp experiments were constrained by the heterogeneity of the presynaptic inputs. CA3 pyramidal neurons are known to project bilaterally to innervate CA1 neurons in both hemispheres (32); in other words, CA1 pyramidal neurons may receive both intra- and inter-hemispherical CA3 inputs. In the EPSC silencing experiment (Fig. 5, D and E), each recorded neuron would receive presynaptic inputs from a mixture of light-sensitive and light-insensitive axons due to unilateral virus transduction. Moreover, expression of piKCR.AT in CA3 pyramidal neurons was achieved by single bolus injection of the virus, which was insufficient to infect all CA3 neurons in the hippocampus. Considering these factors, therefore, the observed piKCR.AT-mediate inhibition effects were highly remarkable. Although piKCR.AT utilizes neurexin’s axonal trafficking and presynaptic stabilization mechanisms, virally expressing piKCR.AT in CA3 pyramidal neurons did not affect the baseline paired-pulse ratio in the EPSC experiments. This observation indicates that localizing piKCR.AT to the presynaptic terminals does not impact synaptic transmission before illumination. Nevertheless, whether expression of piKCR.AT modifies any aspect of synaptic function will require more detailed and comprehensive analysis in the future. The axonal distribution of piKCR.AT in different neuronal projections might vary, and therefore careful evaluations are required before applying the tool in the neurocircuit of interest.

Tools enabling precise control of neuropathways are desirable for resolving circuit mechanisms underlying mental, physiological, and behavioral functions. Although optogenetic excitation of axonal projections has been frequently employed, optogenetic suppression of *bona fide* neuropathway/presynaptic activities encounters more challenges, especially in freely behaving animals. Our results show that piKCR.AT may serve as a robust optogenetic silencer for manipulating neuropathway/presynaptic activities *in vivo*. Cerebellar PCs send axonal projections primarily to the DCN, but they also innervate other regions such as the cerebellar granule layer (42) and the brain stem (43). Previous studies investigated the movement kinematics of mice by optogenetically suppressing PC firing (34, 35), which did not differentiate the functional impacts of distinct PC output pathways. By expressing piKCR.AT in PCs and delivering light to one of their output targets, we were able to optically manipulate the PC-to-DCN pathway and disrupt animal movements on the balance beam. This pilot experiment demonstrates that piKCR.AT allows pathway-targeted neuromodulation in behaving animals, expanding the application repertoire of the HcKCR-based silencers. In conclusion, we have substantially addressed major challenges in HcKCR1 gating and axonal trafficking to develop a KCR-based optogenetic neuropathway silencer (piKCR.AT). The structure-guided mutagenesis study provides insights into the gating mechanism of HcKCR1, and the engineering strategies for piKCR and piKCR.AT may be further employed to discover new generations of KCR tools for neuroscience or biomedical applications.

## Materials and Methods

### Materials

Unless otherwise indicated, chemicals, reagents and buffers were obtained from Sigma-Aldrich (St. Louis, MO, USA) or Thermo Fisher Scientific (Waltham, MA, USA). Light intensity was measured using a S170C microscope slide power sensor (Thorlabs, Newton, NJ, USA) connected to PM100D (Thorlabs) power and energy-meter console.

### Animals

Animal maintenance/handling and experimental protocols were conducted in accordance with the guidelines and regulations of the Institutional Animal Care and Use Committee at Academia Sinica or National Taiwan University. Every effort was made to minimize animal suffering and the number of used animals. Animals were housed at a constant temperature, humidity, and light/dark cycle with *ad libitum* access to food and water. Animals underwent daily monitoring to ensure their gross health status was appropriate. The strain and source of the animals are indicated in the respective experimental sections.

### Cloning and virus preparation

The plasmid of pAAV-CAG-HcKCR1-EYFP was a gift from Professor Mingshan Xue (Addgene plasmid # 182021; http://n2t.net/addgene:182021; RRID: Addgene_182021). Mutations and addition of the trafficking motifs to the HcKCR1 gene were performed using the TOOLS Ultrafast PCR Cloning kit (New Taipei City, Taiwan) and reagents from New England Biolabs (Ipswich, MA, USA). The membrane targeting motif (TS; amino acid sequence: KSRITSEGEYIPLDQIDINV) and ER export motif (EE; amino acid sequence: FCYENEV) were all derived from Kir 2.1 channels. The constructs described in the text were assembled into pAAV vectors containing a CAG promoter (from pAAV-CAG-HcKCR1-EYFP), a CaMKIIα promoter (from pAAV-CaMKIIa-mScarlet; a gift from Professor Karl Diesseroth; Addgene plasmid # 131000; http://n2t.net/addgene:131000; RRID: Addgene_131000), or an EF1α promoter (from pAAV-EF1α1.1-tdTomato; a gift from Professor Edward Boyden; Addgene plasmid # 122101; http://n2t.net/addgene:122101; RRID: Addgene_122101). The DNA fragment of the rat neurexin-1β intracellular domain (amino acid residues 414-468) for constructing piKCR.AT was amplified by PCR from paavCAG-pre-mGRASP-mCerulean (a gift from Professor Jinhyun Kim; Addgene plasmid # 34910; http://n2t.net/addgene:34910; RRID: Addgene_34910). The cloning primers for HcKCR1 mutagenesis are listed in Table S2. Plasmids were amplified in DH5α competent cells (Protech Technology Enterprise Co., Ltd., Taipei, Taiwan) and were purified by HiSpeed Plasmid Midi or Maxi Kits (QIAGEN, Hilden, Germany) for cell transfection or adeno-associated virus (AAV) production. AAV9-CaMKIIa-piKCR.TE, AAV9-CaMKIIα-piKCR.AT, and AAV9-EF1α-piKCR.AT were produced and purified by the Adeno-associated Virus Core Facility of Academia Sinica.

### Expression of the wild-type or engineered HcKCR1 variants in HEK Cells

Human embryonic kidney cells (HEK cells; Elabscience® CL-0001; RRID: CVCL_0045) were first grown in 6-well culture plates and then transfected with plasmid DNAs using Lipofectamine™ 3000 (Invitrogen). After 6–12 h, cells were trypsinized, seeded onto poly-L-lysine coated 12-mm coverslips at 2.5–4 × 10^4^ cells/well in a 24-well culture plate, and then maintained at 37°C and 5% CO_2_. The cell culture medium contained 10% fetal bovine serum (HyClone, Logan, UT, USA) in high glucose Dulbecco’s Modified Eagle Medium supplemented with sodium pyruvate and L-glutamine (Gibco, Thermo Fisher Scientific, Waltham, MA, USA). Electrophysiology recordings were carried out 1–2 days thereafter.

### Electrophysiology (HEK cells)

Whole-cell voltage-clamp recordings were carried out on an IX73 inverted microscope (Olympus, Tokyo, Japan) at room temperature using pipettes pulled from filamented borosilicate capillary glass (Warner Instruments, Hamden, USA) with 3.5–5 MΩ resistance. Cells were held at –60 mV. For Fig. 1, the extracellular solution contained (in mM): 130 NaCl, 2 MgCl_2_, 2 CaCl_2_, 10 HEPES, and 10 Glucose; pH 7.4. The intracellular solution contained (in mM): 130 KCl, 2 MgCl_2_, and 10 HEPES; pH 7.2. For Fig. 2, the extracellular solution contained (in mM): 138 NaCl, 1.5 KCl, 1.2 MgCl_2_, 2.5 CaCl_2_, 5 HEPES, and 10 Glucose; pH 7.4. The intracellular solution contained (in mM): 135 K-gluconate, 10 NaCl, 2 MgCl_2_, 10 HEPES, 2 Mg ATP, 1 EGTA; pH 7.2. Signals were amplified using a MultiClamp 700B microelectrode amplifier (Molecular Devices, San Jose, CA, USA), low-pass filtered at 2 kHz, digitized at 10 kHz by a Digidata 1550B acquisition system (Molecular Devices), and recorded with software pClamp 10 (Molecular Devices). Light for photocontrol was provided by a pE-4000 illumination system (CoolLED, Andover, UK), and was projected to the recorded cell through a 40X microscope objective.

### Preparation and culture of primary cortical neurons

Pregnant wild-type Sprague-Dawley rats (from BioLASCO Taiwan Co., Ltd., Taipei, Taiwan) were anesthetized by inhalation of isoflurane (2-3%), followed by cervical dislocation. Brains from E18–19 embryos were harvested and dissected to isolate the cortical tissues. After papain digestion (17.5 U/mL) and gentle trituration, the cells were seeded onto 12-mm coverslips (coated with poly-L-lysine; 0.5 mg/mL in 50 mM borate buffer) at 8 × 10^4^ cells/well in a 24-well culture plate. The cells were initially cultured at 37°C and 5% CO_2_ in the neuron culture medium (NCM) supplemented with 5% fetal bovine serum (HyClone, Logan, UT, USA). The NCM included Neurobasal Medium (Gibco, Thermo Fisher Scientific, Waltham, MA, USA), 2% B-27™ Supplement (Gibco), 1% Penicillin-Streptomycin (Gibco), and 0.25% GlutaMAX (Gibco). After 6–8 h, the medium was replaced by serum-free NCM. The cells were maintained at 37°C and 5% CO_2_, and half of the medium was replaced with fresh serum-free NCM every week.

### Expression of piKCR.TE or HcKCR1.TE in cultured neurons

Primary cortical neurons (10–12 DIV) were transfected with the DNA of piKCR.TE or HcKCR1.TE (in pAAV-CaMKIIa vector; 0.25 μg/coverslip) using Lipofectamine^TM^ 2000 (Invitrogen). Prior to transfection, half of the cultured medium was taken out for the preparation of the conditioned medium (50% original and 50% fresh). To ensure proper DNA packaging by lipofectamine without overly expressing the opsins (which may cause neuronal toxicity), neurons were co-transfected with an empty vector pCAGEN (0.75 μg/coverslip). After incubating cells with the DNA-lipofectamine mixture for 30 minutes, the medium was replaced with the conditioned medium for cell maintenance. Electrophysiology recordings were carried out 2–4 days thereafter.

### Electrophysiology (cultured neurons)

Whole-cell current-clamp recordings were carried out in 13–21 DIV cortical neurons using the same devices for HEK cell recordings. Pipette resistance was 4–6.5 MΩ. Neurons were held at – 60 mV. The extracellular solution contained (in mM): 138 NaCl, 1.5 KCl, 1.2 MgCl_2_, 2.5 CaCl_2_, 5 HEPES, 10 Glucose; pH 7.4. The intracellular solution contained (in mM): 135 K-gluconate, 10 NaCl, 2 MgCl_2_, 10 HEPES, 2 Mg ATP, 1 EGTA; pH 7.2. Signals were amplified using a MultiClamp 700B microelectrode amplifier (Molecular Devices), low-pass filtered at 2 kHz, digitized at 10 kHz by a Digidata 1550B acquisition system (Molecular Devices), and recorded with software pClamp 10 (Molecular Devices). Light for photocontrol was provided by a pE-4000 illumination system (CoolLED) and was projected to the recorded cell through the 40X microscope objective.

### Confocal microscopy (HEK cells and cultured neurons)

To compare the expression pattern of piKCR vs. piKCR.TE (Fig. 2A), HEK cells were rinsed with Phosphate Buffered Saline (PBS) followed by fixation in 4% paraformaldehyde (PFA; in PBS) at room temperature for 10 min. To check neuronal expression of piKCR.TE and HcKCR1.TE (Fig. 3A), cultured neurons were rinsed with warm (37°C) Dulbecco’s Phosphate-Buffered Saline (DPBS) supplemented with 4% sucrose, followed by fixation in warm 4% PFA + 4% sucrose for 10 min. The fixed cells were then washed with PBS and mounted in Anti-Fade Fluorescence Mounting Medium (Abcam ab104135). Confocal images were acquired using a 63X oil-immersion objective on an LSM 880 confocal microscope (Zeiss, Jena, Germany).

### Expression of piKCR.TE in the anterior paraventricular nucleus of the thalamus (PVA)

Wild-type C57B6 mice (8–12 weeks old) were fixed in a stereotaxic head frame after being anesthetized with 2% isoflurane. The skull was made visible by shaving the scalp and making a midline incision. The cranial fissures were used to align the skull. After making the hole with a drill bit (MA Ford, #87), a pulled glass capillary (tip diameter ∼20–40 μm) backfilled with the viral solution (AAV9-CaMKIIα-piKCR.TE, 1.2 × 10^13^ vg/mL) was introduced via the hole to reach the target area. Using a Hamilton micro-syringe and computer-assisted automatically controlled pump, 200 nL of the virus was administered (AP 0.34 mm, ML 0.00 mm, DV –3.5 mm) at 40 nL/min. After being in place for 5–10 min, the injection needle was progressively removed. After four weeks of recovery, mice were subjected to electrophysiology experiments as described below.

### Electrophysiology (PVA neurons)

Animals expressing piKCR.TE in the PVA neurons were deeply anesthetized with isoflurane and decapitated after confirming the absence of nocifensive responses. Briefly, brains were rapidly removed and transferred into oxygenated cutting solution (95% O_2_, 5% CO_2_) containing (in mM): 110 choline chloride, 25 NaHCO_3_, 25 D-glucose, 12.7 ascorbate, 7 MgSO_4_, 3.1 Na-pyruvate, 2.5 KCl, 1.25 NaH_2_PO_4_, and 0.5 CaCl_2_. Coronal slices (300 μm) containing the PVA were prepared on a vibratome (5100mz-Plus, Campden Instruments, Loughborough, UK), and incubated in a holding chamber for 60 min at 32 °C to allow recovery. The recovered slices were transferred to the recording chamber of a SliceScope Pro 6000 (Scientifica, UK) and continuously perfused (2–4 mL/min) with oxygenated artificial cerebrospinal fluid (aCSF) containing (in mM): 119 NaCl, 26.2 NaHCO_3_, 11 glucose, 2.5 KCl, 2 CaCl_2_, 1.3 MgSO_4_, and 1 NaH_2_PO_4_. Neurons were visualized using a 40X water-immersion objective on an upright fluorescence microscope (BX51, Olympus) equipped with infrared differential interference contrast (IR-DIC) optics and epifluorescence. Whole-cell current clamp recordings were carried out using pipettes pulled from borosilicate glass capillaries (Warner Instruments). The patch pipettes were pulled by an MP-500 micropipette puller (RWD Life Science, Shenzhen, China) to yield a tip resistance of 5–6.5 MΩ. The pipette internal solution contained (in mM): 131 K-gluconate, 20 KCl, 4 NaCl, 2 EGTA, 10 HEPES, 2 Na-ATP, and 0.3 Na-GTP (pH 7.3, 290 mOsm). Membrane potentials were recorded with a Multiclamp 700B amplifier (Molecular Devices). Signals were sampled at 50 kHz and low-pass filtered at 2 kHz. Voltage and current commands were generated and digitized with a Digidata 1440A, controlled by Clampex 11.2 software (Molecular Devices). An LED light source (CoolLED pE-300white, Andover, UK) was used to optically suppress action potential generation. Yellow light was generated by filtering white light through a TEXRED cube set (Ex 560/40 nm, Em 590 lp, Dichroic T590lpxr; Chroma 49017), and was delivered to the recorded neurons through the microscope objective. All recordings were performed at room temperature (27–29 °C).

### Expression of piKCR.TE, piKCR.AT, or EYFP in the hippocampus

C57BL/6 mice (8‒10 weeks old; BioLASCO Taiwan Co., Ltd.) were anesthetized using isoflurane (induction: 5%; during surgery: 1‒2%). Each anesthetized mouse was placed in a stereotaxic frame (RWD Life Science, Shenzhen, China) for surgery. The eyes of the mouse were protected with Vaseline. The head fur of the mouse was trimmed close to skin, and the trimmed region was cleaned with 70% ethanol. A midline incision of the scalp was carried out to expose the skull. The scalp was disinfected by povidone iodine, followed by mild tissue clearing with 3% hydrogen peroxide. A small hole was drilled into the skull bone for the targeted brain region. The mouse was injected unilaterally with 200 nL of virus suspension at a rate of 50 nL/min using a Hamilton 80000 Gastight Syringe (Hamilton Company, Inc., NV, USA) attached to a syringe pump. The following coordinates were used for targeting the hippocampal CA3 region: anteroposterior (AP) –2.45, mediolateral (ML) –2.35, and dorsoventral (DV) –3.35. For confocal imaging experiments in Fig. 5, the injected virus suspension contained either AAV9-CaMKIIα-piKCR.TE or AAV9-CaMKIIα-piKCR.AT (1.2 × 10^9^ vg/μL) mixed with AAV9-CaMKIIα-mScarlet (as an injection marker; 5 × 10^7^ vg/μL). For electrophysiology experiments in Fig. 5/Fig. S4 and Fig. S5, ∼1 × 10^9^ vg/μL of AAV9-CaMKIIα-piKCR.AT and AAV9-CaMKIIα-EYFP was injected, respectively. After injection was completed, the microsyringe was left in place for an additional 10 minutes to allow viral diffusion. The microsyringe was then slowly withdrawn from the brain, followed by skin closure using sterile sutures. To prevent infection and reduce discomfort of the animal, the mouse was subcutaneously injected with ampicillin (24 mg/kg) and Carprofen (5 mg/kg). After 2–4 weeks of recovery, mice were subjected to imaging or electrophysiology experiments as described below.

### Preparation of brain sections for confocal imaging (hippocampus and PVA)

Mice expressing piKCR.TE or piKCR.AT in the hippocampus (10–12 weeks old, i.e., 2–4 weeks after virus injection) were deeply anesthetized and perfused with PBS, followed by an infusion of 4% PFA (in PBS). Brains were collected and post-fixed overnight in 4% PFA at 4°C, washed the next day by PBS, and then sectioned using a DTK-1000N MicroSlicer (Dosaka, Kyoto, Japan) to obtain coronal sections at 300-μm thickness. The brain slices were further cleared by RapiClear^®^ 1.49 (SunJin Lab Co., Hsinchu, Taiwan) following manufacturer’s instructions. Near-horizontal hippocampus-containing brain sections (used for electrophysiology experiments in Fig. 5, S4, and S5) and coronal PVA-containing brain sections (used for electrophysiology experiments in Fig. 4) were also postfixed by 4% PFA at 4°C overnight and treated with RapiClear^®^ 1.49 prior to imaging acquisition.

### Confocal image acquisition and analysis (hippocampus)

Tiled images of the cleared brain slices (mounted in RapiClear^®^ 1.49) were acquired using a 20X objective on an LSM 880 (for coronal sections) or LSM 700 Stage (for near-horizontal sections) confocal microscope (Zeiss, Jena, Germany). For coronal sections, the EYFP fluorescence signal from piKCR.TE or piKCR.AT was analyzed using the MetaMorph software (Molecular Devices). In each image, regions of interest (ROIs, 100 × 40 pixels) were selected in the ipsilateral CA3 cell-body layer, contralateral CA1 SO, and contralateral CA1 SR (three ROIs in each location, together with a background ROI in the nearby uninfected area). For each location, fluorescence densities of the three selected ROIs were first background-subtracted and then averaged. The average fluorescence densities of contralateral CA1 SO and contralateral CA1 SR were individually normalized by the average fluorescence density of ipsilateral CA3 cell-body layer in the same image. The “normalized fluorescence densities” of the contralateral CA1 sublayers were used to compare the axon-targeting effects of piKCR.TE or piKCR.AT (Fig. 5C).

### Electrophysiology (hippocampal neurons)

Hippocampal slices were prepared from virus-injected mice described above. In brief, mice were decapitated under deep isoflurane anesthesia, and their brains were quickly removed and transferred to an ice-cold dissection solution containing (in mM): 204.5 sucrose, 2.5 KCl, 1.25 NaH_2_PO_4_, 28 NaHCO_3_, 7 dextrose, 3 Na-pyruvate, 1 Na-ascorbate, 7 MgCl_2_ (pH 7.3, osmolarity 310–320, oxygenated with 95% CO_2_ and 5% O_2_). 350-μm-thick slices were sectioned at oblique angles using a vibroslicer (VT 1200S, Leica Microsystems, Wetzlar, Germany). An incision was made to separate the CA1 and CA3 regions to suppress epileptiform activity in CA1. Before recording, slices were incubated for 30 min at 35–37°C in standard artificial cerebrospinal fluid (aCSF) containing (in mM): 125 NaCl, 2.5 KCl, 1.25 NaH_2_PO_4_, 25 NaHCO_3_, 25 dextrose, 2 CaCl_2_, 1 MgCl_2_, 3 Na-pyruvate, 1 Na-ascorbate (pH 7.3, osmolarity 295–300, oxygenated with 95% CO_2_ and 5% O_2_). The recovery chamber was then maintained for at least 1 h at room temperature (25–27°C) prior to recording.

Hippocampal CA1 pyramidal neurons were visually identified using a 40x water immersion objective (Olympus) equipped with an IR-DIC (infrared-differential interference contrast) upright microscope (BX51WI, Olympus). Slices were constantly perfused with oxygenated aCSF at 31–33°C. SR-95531 (gabazine; 1–2 μM) was routinely added to block GABA_A_ receptors. Whole-cell recordings were carried out using an EPC-10 patch-clamp amplifier controlled by PatchMaster software (HEKA, Lambrecht (Pfalz), Germany) after online low-pass filtering at 5 kHz and digitizing at 50 kHz. CA1 pyramidal neurons were voltage-clamped at a holding potential of –70 mV. The recording pipette was pulled to a resistance of 6–8 MΩ when filled with an intracellular solution containing (in mM): 130 K-gluconate, 10 KCl, 10 Na_2_-phosphocreatin, 10 HEPES, 4 Mg-ATP, 0.3 Na-GTP (pH 7.3, osmolarity 295–300). Series resistance (15–20 MΩ) was continuously monitored during recording. Cells were discarded if series resistance was over 20 MΩ and/or with a drift >20%.

Excitatory postsynaptic currents (EPSCs) were evoked by paired-pulse stimulation (0.1 ms in duration and 50 ms in inter-pulse interval) with 0.05 Hz of each epoch, using a bipolar tungsten electrode (FHC, Bowdoin, ME, USA) positioned on the *stratum radiatum* (SR) ∼200–300 μm away (toward subiculum) from the recorded cell to electrically recruit synaptic inputs from the CA3 neurons. In both dark and light conditions, EPSCs were recorded for 10 epochs. A pE-300 light engine (CoolLED) was used for piKCR photostimulation. To test whether piKCR.AT enables EPSC inhibition presynaptically, a flash of 560 nm light (0.93 mW/mm^2^, started 50 ms prior to the first stimulation and ended after the second pulse) was delivered through the microscope objective to cover the duration of paired-pulse stimulation.

### Optical manipulation of the PC-to-DCN pathway in behaving mice

*Surgical procedures.* Male and female C57BL/6J mice (Jackson Laboratory #000664) between 6 to 8 months of age were used. AAV9 encoding piKCR-EYFP.AT (under the control of EF1α promoter) was stereotaxically injected into the cerebellar cortex of each mouse at two locations (AP, –6.24 mm; ML, 0 mm; DV, –0.3 and –0.8 mm from dura). Each location received 2.5 μL of virus (8.6 × 10^11^ vg/mL) at a speed of 0.2 μL/min. The mice were trained/tested both before and 14 days after viral injection to ensure that their behaviors were not affected by the surgical procedures. Three weeks after virus injection, a homemade optrode was implanted in the deep cerebellar nuclei of each mouse (AP, –6.24 mm; ML, ±2.1 mm; DV, –1.9 mm from dura). The optical control experiments were carried out 4–6 weeks after virus injection.

*Behavioral procedures.* A balance beam walking test was performed to assess the capability of animals to travel across a beam of 1-cm wide and 100-cm long (suspended 12 cm above the bench) for reaching a cage. The mouse was placed before the start line and was removed from the beam once it had crossed the finishing line. To reduce the stress elicited by unfamiliar experimental procedures, mice were allowed to adapt in the room for 15 minutes prior to the test each day. The experiment was preceded by 4 days of habituation. The test was then repeated for over 3 consecutive days, with mice being able to cross the apparatus at least two times each day. Trials were separated by a 5-minute break. If the mouse went in the opposite direction, the trial was considered as incomplete. To inhibit the axonal terminals of Purkinje cells, pulses of 589-nm light (Changchun New Industries Optoelectronics Tech. Co.,Ltd., China; 27–30 mW measured at the output of light source) was delivered at 40 Hz with 50% duty cycle. The illumination was calibrated by a power meter (Thorlabs) to ensure consistent intensity throughout the experiment. Mann-Whitney U test was carried out with a two-tailed P value to assess group differences.

### Confocal imaging of the cerebellar slices

Mice that underwent behavioral tests were perfused (first with PBS, followed by 4% PFA in PBS) for brain collection. The collected brains were post-fixed in 4% PFA (in PBS) and washed by PBS prior to sectioning. Sixty-μm-thick sagittal sections were obtained using a vibratome machine (Model 5100mz, Campden Instruments, UK), treated with DAPI (4’,6-diamidino-2-phenylindole) for nuclear counterstaining, and then mounted on glass slides in VECTASHIELD Antifade Mounting Medium (Vector Laboratories, CA, USA). Tiled images of the entire cerebellar cross-section (Fig. 6B, left) and the DCN (Fig.e 6B, right) were acquired using a 20X and a 40X objective lens on an LSM 700 Stage confocal microscope (Zeiss).

### Data Analysis

The electrophysiology data from HEK cells and cultured neurons were analyzed in Clampfit 10 (Molecular Devices). The electrophysiology data from PVA neurons were analyzed with pCLAMP 11.2 (Molecular Devices). The P_K_/P_Na_ permeability ratio for the wild-type and engineered HcKCR1 variants was calculated using the modified Goldman-Hodgkin-Katz (GHK) voltage-equation. The decay time constant from Fig. 1F was calculated by a single exponential fit. Data plotting and statistical analysis were carried out using GraphPad Prism (Dotmatics, Boston, MA, USA). The sample size (n), *P* value range, and statistical test performed for each experiment are included in the respective figure legend or main text.

## Acknowledgments

We are grateful for the professional supports provided by Academia Sinica DNA Sequencing Core Facility, Academia Sinica AAV Core Facility, and the Light Microscopy Core Facility of the Institute of Biomedical Sciences, Academia Sinica. We also thank Mr. Yan-Ting Guo, Mr. Ching-Chuan Cheng and Ms. Yu-Yen Lee for assisting in cloning, electrophysiology and confocal microscopy experiments.

## Author contributions

Conceptualization: W.C.L., S.M.M.L., C.L.H., M.K.P., C.C.C.

KCR engineering: S.M.M.L.

Electrophysiology (HEK cells and cultured neurons): S.M.M.L.

PVA experiments (virus injection and electrophysiology): W.H.C. and Y.C.C.

Hippocampus experiments (virus injection, imaging, and electrophysiology): H.Y.W. and Y.J.L.

Behavior experiment: I.C.L. Visualization: W.C.L. and S.M.M.L.

Supervision: W.C.L., C.L.H., M.K.P., C.C.C.

Writing: W.C.L. and S.M.M.L., with contribution from all authors.

## Data and materials availability

All data needed for the conclusions of this study are presented in the paper and/or the Supplementary Materials.

