## Supplemental Materials for "Engineering a performance-improved, axon-targeted kalium channelrhodopsin for optogenetic neuropathway inhibition"

Simon Miguel M. Lopez *et al.*

**This PDF file includes:**

Figures S1 to S5

Tables S1 and S2

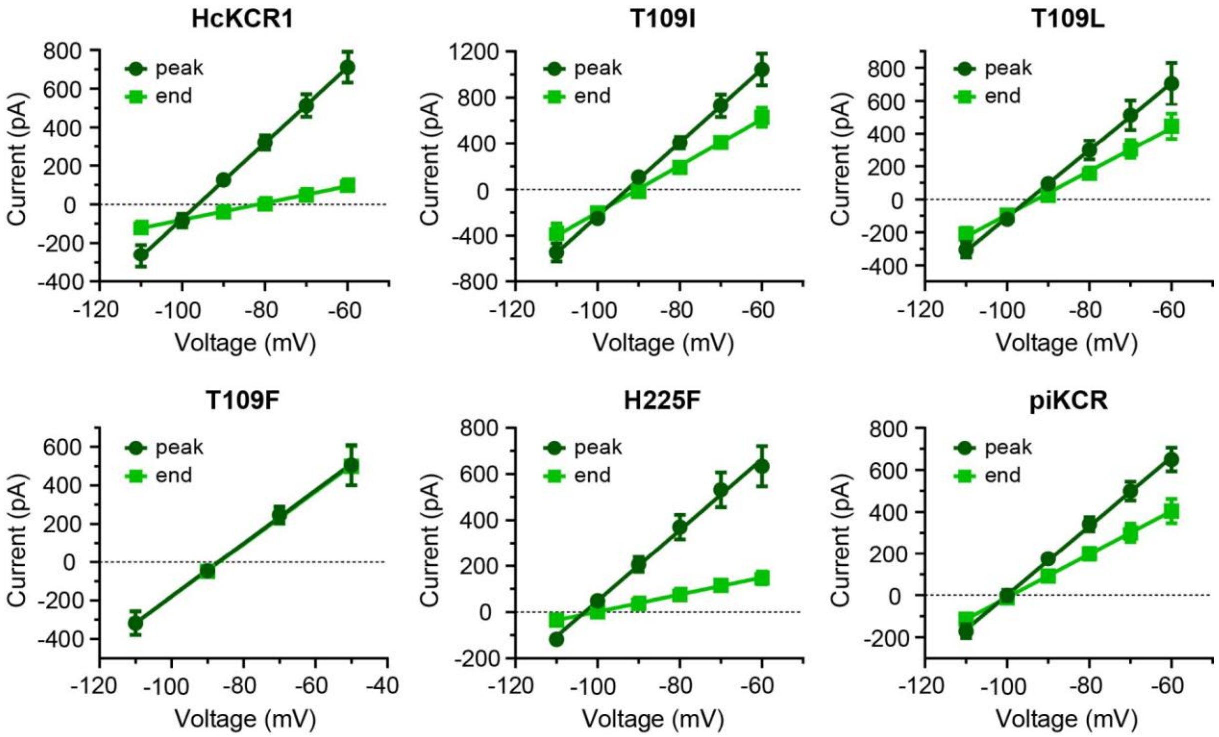


**Fig. S1. Current-voltage relations (I-V curves) of the HcKCR1 variants shown in Fig. 1H.** Opsins were expressed in HEK cells, and currents were elicited by 500 ms of 550 nm illumination (0.084 mW/mm$^2$) under the ionic conditions shown in Fig. 1B. Current amplitudes were measured at the peak (dark green) and at the end of the illumination (light green). Data are plotted as mean ± SEM. n = 9 cells for HcKCR1, T109I, and T109L. n = 8 cells for T109F, KALI-1, and piKCR.

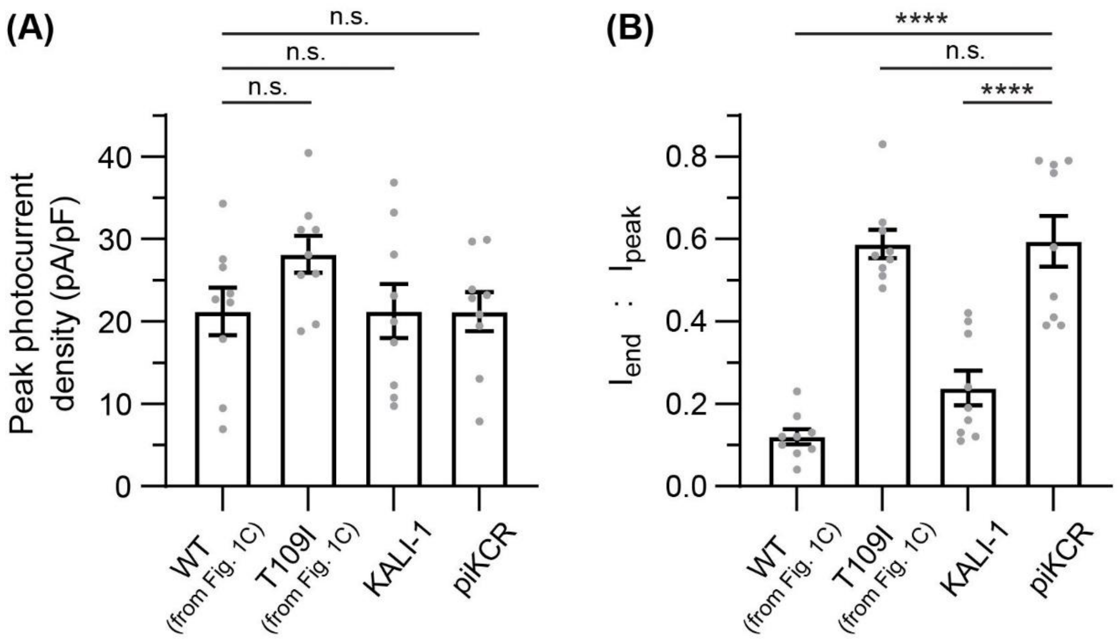


**Fig. S2. Photocurrent density (A) and end-to-peak ratio (B) of the wild-type HcKCR1, the T109I mutant, the H225F mutant (KALI-1), and the T109I+H225F double mutant (piKCR).** piKCR elicited comparable photocurrent magnitude with the wild-type (**A**) and exhibited similarly slow desensitization (manifested as the high end-to-peak current ratio) as the T109I mutant (**B**). Opsins were expressed in HEK cells, and currents were elicited by 500 ms of 550 nm illumination (0.084 mW/mm$^2$) under the ionic conditions shown in Fig. **1B**. Cells were held at –60 mV. $I_{end}$: photocurrent amplitude measured at the end of the illumination. $I_{peak}$: amplitude of the photocurrent peak. Data are plotted as mean ± SEM, with individual measurements shown as grey symbols. n = 9 cells for all tested opsins. ****$P < 0.0001$; n.s., not significant; one-way ANOVA.

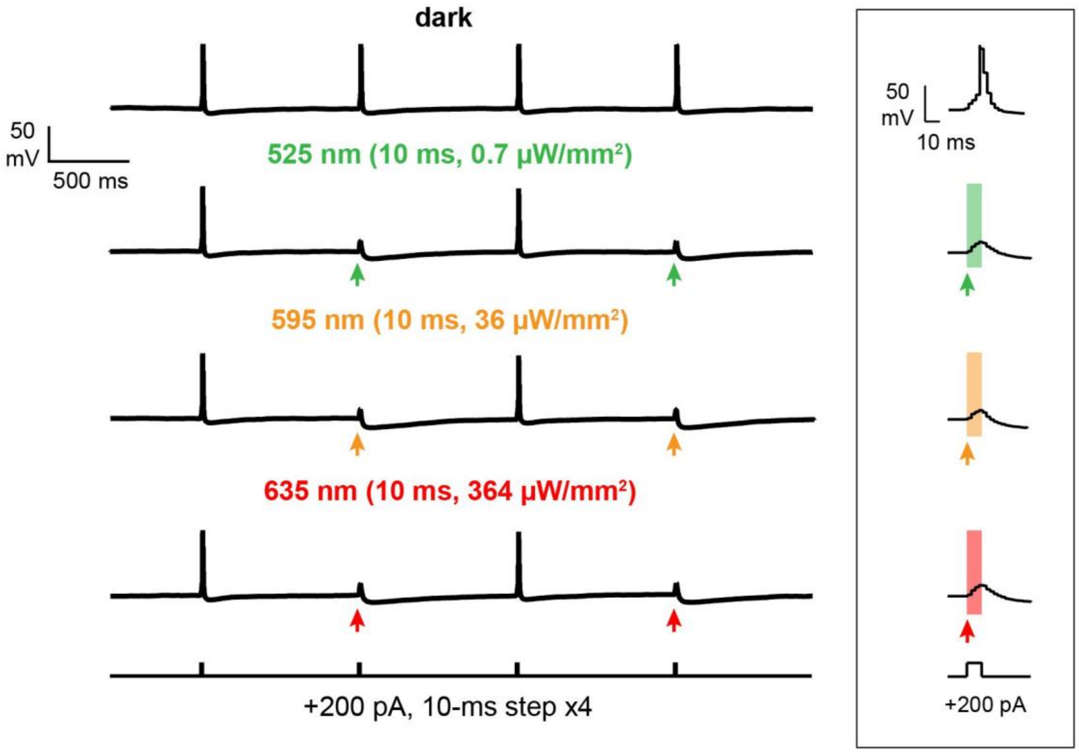


**Fig. S3. piKCR.TE can achieve instantaneous optical suppression of neuronal spiking.** Whole-cell current clamp recordings from a cultured cortical neuron expressing piKCR.TE. Single action potentials were elicited every second by a 10-ms current step (+200 pA). Each colored arrow indicates a 10-ms light flash that was delivered simultaneously with a current injection. Using the light intensities indicated above, the 10-ms flashes were sufficient to prevent action potential firing and allowed rapid recovery of neuronal excitability (i.e., successful spike generation by the succeeding current injection).

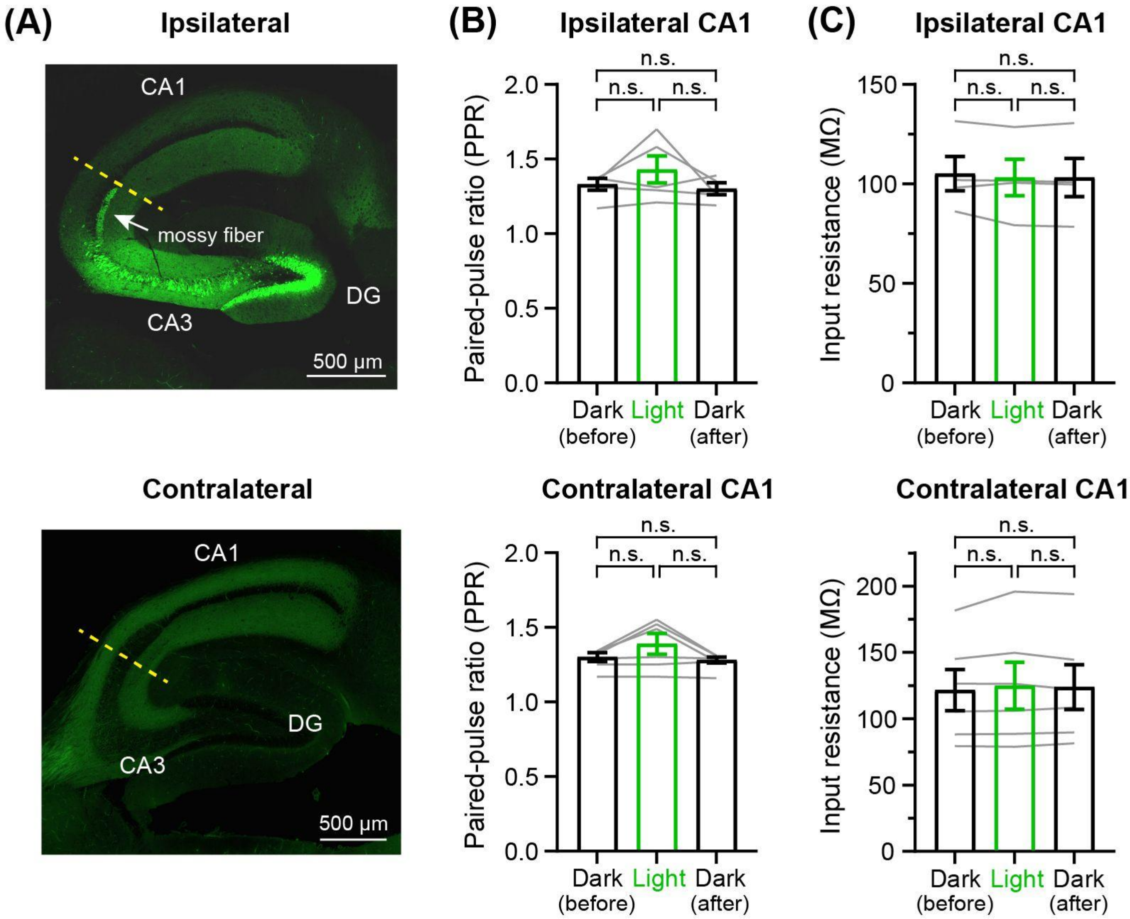


**Fig. S4. Photostimulation of piKCR.AT in the CA3→CA1 pathways does not significantly alter paired pulse ratio (PPR) or input resistance of the recorded cells.** (**A**) Confocal images of near-horizontal hippocampal slices for EPSC recordings (Fig. 5, panels **D** and **E**). The green color indicates the distribution of piKCR.AT. Slices were prepared from mice unilaterally injected with AAV9 encoding piKCR.AT. An incision was made in each recorded slice (indicated by yellow dashed lines) to exclude somatic effects of the opsin in CA3 neurons (ipsilateral slice) and prevent epileptiform activity in CA1. The fluorescent cell bodies in the ipsilateral slice indicate the infected regions. In this example, not only CA3 but also DG were infected by the virus. Thus, EYFP fluorescence was detected in both CA1 (innervated by CA3 pyramidal neurons) and the mossy fiber (which consists of axonal projections from the DG granule cells). (**B**) PPR from the recorded cells in Fig. **5E**; mean ± SEM, n = 5 and 6 cells from ipsilateral and contralateral slices, respectively. (C) Input resistance of the recorded cells in Fig. **5E**; mean ± SEM, n = 4 and 6 cells from ipsilateral and contralateral slices, respectively. Measurements from individual cells are plotted as grey lines. n.s., not significant (repeated-measures one-way ANOVA with Geisser–Greenhouse correction followed by Tukey's comparison).

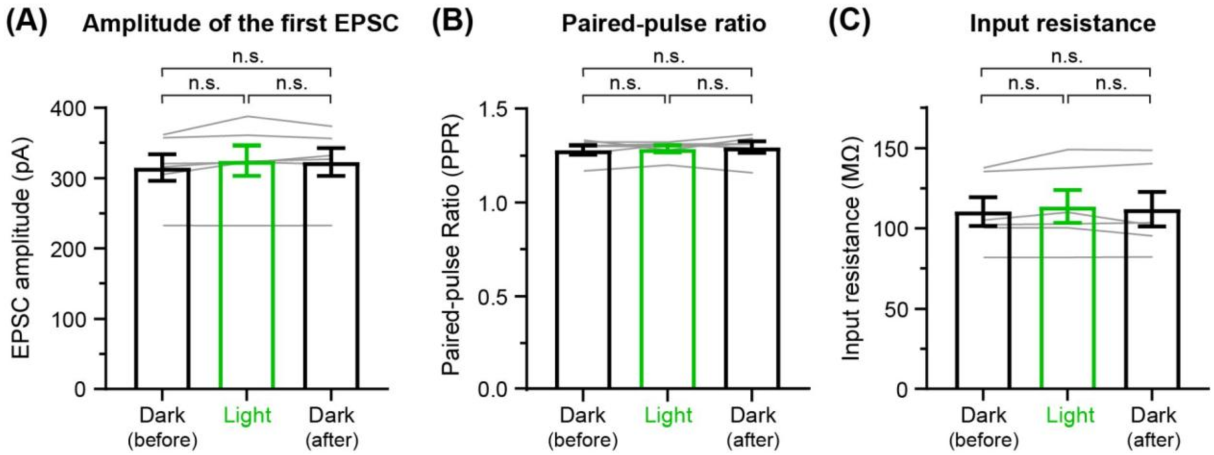


**Fig. S5. Green light does not affect the EPSC amplitude (A), paired-pulse ratio (B), or input resistance (C) of CA1 pyramidal cells downstream of EYFP-expressing CA3 neurons.** EYFP was unilaterally expressed in CA3 pyramidal neurons via stereotaxic injection of AAV9–CaMKIIα–EYFP using the same conditions/procedures as those for AAV9–CaMKIIα–piKCR.AT in Fig. 5 and S4. Ipsilateral CA1 pyramidal neurons were held at –70 mV, and EPSCs were electrically evoked by paired-pulse stimulation in the SR as described in Fig. 5. Data are plotted as mean ± SEM. Measurements from individual cells are plotted as grey lines. n = 6 cells. n.s., not significant (repeated-measures one-way ANOVA with Geisser–Greenhouse correction followed by Tukey’s comparison).

**Table S1. Analysis of reversal potentials for HcKCR1 variants in Fig. 1H**

| | | | | ***P*-value (vs. wild-type peak)**[c] | |
|---|---|---|---|---|---|
| **HcKCR1 variant** | **n** | **Reversal potential drift from peak to end (mV)**[a] | ***P*-value (peak vs. end reversal potential)**[b] | **Peak Reversal Potential** | **End Reversal Potential** |
| Wild-type | 9 | 14.9 ± 0.9 | <0.0001 (****) | -- | -- |
| T109I | 9 | 2.1 ± 0.2 | <0.0001 (****) | 0.0183 (*) | 0.0005 (***) |
| T109L | 9 | 1.7 ± 0.3 | 0.0003 (***) | 0.2319 (ns) | <0.0001 (****) |
| T109F | 8 | 0.13 ± 0.13 | 0.3579 (ns) | <0.0001 (****) | 0.0801 (ns) |
| KALI-1 | 8 | 0.8 ± 0.6 | 0.1881 (ns) | 0.0015 (**) | <0.0001 (****) |
| piKCR | 8 | 1.0 ± 0.2 | 0.0033 (**) | 0.1522 (ns) | <0.0001 (****) |

[a] Mean ± SEM.
[b] *$P$ < 0.05; **$P$ < 0.01; ***$P$ < 0.001; ****$P$ < 0.0001. Two-tailed paired t test.
[c] Comparison with the peak reversal potential of the wild-type. *$P$ < 0.05; **$P$ < 0.01; ***$P$ < 0.001; ****$P$ < 0.0001. ns, not significant. One-way ANOVA with Dunnett's multiple comparison test.

**Table S2. List of primers used for HcKCR1 mutagenesis**

| HcKCR1 Variant | DNA Fragment # | Forward Sequence (5'→3') | Reverse Sequence (5'→3') |
|---|---|---|---|
| T109A | 1 | gaattgcggcccaacggtaccatgcctttctacgac | gatcagcagggggcatgcgaacacgtagtccag |
| | 2 | ctggactacgtgttcgcatgccccctgctgatc | ctcgcccttgctcacgttaacggacagtgtctcag |
| T109V | 1 | gaattgcggcccaacggtaccatgcctttctacgac | gatcagcagggggcagacgaacacgtagtccag |
| | 2 | ctggactacgtgttcgtctgccccctgctgatc | ctcgcccttgctcacgttaacggacagtgtctcag |
| T109I | 1 | gaattgcggcccaacggtaccatgcctttctacgac | ggatcagcagggggcagatgaacacgtagtccag |
| | 2 | ctggactacgtgttcatctgccccctgctgatcc | ctcgcccttgctcacgttaacggacagtgtctcag |
| T109L | 1 | gaattgcggcccaacggtaccatgcctttctacgac | gatcagcagggggcacaggaacacgtagtccag |
| | 2 | ctggactacgtgttcctgtgccccctgctgatc | ctcgcccttgctcacgttaacggacagtgtctcag |
| T109N | 1 | gaattgcggcccaacggtaccatgcctttctacgac | ggatcagcagggggcagttgaacacgtagtccag |
| | 2 | ctggactacgtgttcaactgccccctgctgatcc | ctcgcccttgctcacgttaacggacagtgtctcag |
| T109M | 1 | gaattgcggcccaacggtaccatgcctttctacgac | gatcagcagggggcacatgaacacgtagtccag |
| | 2 | ctggactacgtgttcatgtgccccctgctgatc | ctcgcccttgctcacgttaacggacagtgtctcag |
| T109F | 1 | gaattgcggcccaacggtaccatgcctttctacgac | gatcagcagggggcagaagaacacgtagtccag |
| | 2 | ctggactacgtgttcttctgccccctgctgatc | ctcgcccttgctcacgttaacggacagtgtctcag |
| T109H | 1 | gaattgcggcccaacggtaccatgcctttctacgac | ggatcagcagggggcaatggaacacgtagtccag |
| | 2 | ctggactacgtgttccattgccccctgctgatcc | ctcgcccttgctcacgttaacggacagtgtctcag |
| H225F | 1 | gaattgcggcccaacggtaccatgcctttctacgac | gatccagaaaggcgaagatgatgtagaagg |
| | 2 | ccttctacatcatcttcgcctttctggatc | ctcgcccttgctcacgttaacggacagtgtctcag |
